# Targeted disruption of phage liquid crystalline droplets abolishes antibiotic tolerance of bacterial biofilms

**DOI:** 10.1101/2025.07.28.667140

**Authors:** Abul K. Tarafder, Miles Graham, Luke K. Davis, Shawna Pratt, Jan Böhning, Pavithra Manivannan, Zhexin Wang, Camila M. Clemente, Raymond J Owens, George A. O’Toole, Philip Pearce, Tanmay A.M. Bharat

## Abstract

All bacterial biofilms contain an extracellular matrix rich in filamentous molecules ^1^ that self-associate ^2–4^, conferring emergent properties to bacteria ^5^, including antibiotic tolerance ^6^. *Pseudomonas aeruginosa* is a human pathogen that forms biofilms in diverse infectious settings ^7,8^, where the upregulation of a filamentous bacteriophage Pf4, has been shown to be a key virulence factor that protects bacteria from antibiotics ^9,10^. Here, we modelled biophysical characteristics of biofilm-linked liquid crystalline droplets formed by Pf4, which predicted that sub-stoichiometric phage binders had the ability to disrupt liquid crystals by changing the surface properties of the phage. We tested this prediction by developing nanobodies targeting the outer surface of the Pf4 phage, which disrupted *in vitro* reconstituted droplets, promoted antibiotic diffusion into bacteria, disrupted *P. aeruginosa* biofilm formation under a variety of conditions, and abolished antibiotic tolerance of biofilms. The inhibition strategy illustrated in this study could be extended to biofilms of other pathogenic bacteria, where filamentous molecules are pervasive in the extracellular matrix. Furthermore, our findings exemplify how targeting a biophysical mechanism, rather than a defined biochemical target, is a promising avenue for intervention, with the potential of applying this concept to other disease-related contexts.

## Introduction

The perennial rise of antibiotic tolerant bacterial infections is a serious threat to human health ^11^, leading to an urgent need to understand fundamental mechanisms of antibiotic tolerance in order to develop new strategies for treating infections ^12^. *Pseudomonas aeruginosa*, an opportunistic human pathogen, is one of the leading causes of human morbidity and mortality ^13^. *P. aeruginosa* commonly infects bones, wounds, lungs (especially in persons with cystic fibrosis) as well as medical hardware in hospital settings ^7,8,14^. The majority of chronic *P. aeruginosa* infections proceed via the formation of biofilms ^15^, which are multicellular communities of bacteria encased in a primarily self-produced extracellular polymeric substance (EPS) matrix. The EPS matrix is composed of biopolymers such as extracellular DNA (eDNA), polysaccharides and filamentous protein fibers ^2-4,16,17^. The EPS matrix is a defining feature of biofilms, which bestows beneficial emergent properties to bacteria, such as immune evasion, the ability to withstand external physical stresses, and up to a thousand-fold increased tolerance to antibiotics ^5,6,18,19^.

The most virulent of *P. aeruginosa* isolates encode an inoviral Pf prophage in their genome ^20,21^. Inoviruses are filamentous, single-stranded DNA-containing phages that can be several micrometres in length and ~6 nm in diameter ^22^. Upon switching to a biofilm lifestyle, the expression of the inovirus Pf4 is highly upregulated by *P. aeruginosa* ^23^ and Pf4 phages secreted from the bacteria form an integral part of the biofilm matrix ^9^. Pf4 is symbiotic with *P. aeruginosa* and has been found in the most virulent clinical isolates, correlating with increased bacterial pathogenicity and morbidity ^24,25^. Pf4 increases bacterial pathogenicity by acting as an immune decoy in wound infection models ^26^, decreasing bacterial phagocytic uptake by macrophages ^27^ and inhibiting wound epithelialization by keratinocytes ^28^.

Pf4 phage filaments also promote biofilm stability by forming spindle-shaped liquid crystalline droplets termed tactoids ^9^. These structures form by depletion attraction interactions, where filamentous phage particles align along their long axis in the presence of biopolymer, which is excluded in the process. The alignment of phages into liquid crystalline droplets permits higher degrees of translational and orientational freedom of the biopolymer, which is entropically favourable. Thus, depletion attraction is a biophysical, entropically driven process, rather than being driven by specific biochemical interactions ^9,29^. Previously we showed that these Pf4 liquid crystalline droplets encapsulate bacterial cells ^10^, forming a protective shield that reduces antibiotic diffusion, thereby increasing antibiotic tolerance ^10,30^. Bacterial encapsulation by liquid crystalline droplets is also driven by biophysical characteristics, governed by phage filament shape and packing rather than specific biochemical interactions between phage or the bacterial cell surface ^30^.

In this study, starting with biophysical modeling that predicted that a sub-stoichiometric binder could disrupt Pf4 liquid crystalline droplets, we developed nanobody inhibitors against Pf4 that not only disrupted Pf4 droplets under various conditions, but also abolished antibiotic tolerance of *P. aeruginosa* biofilms ^31,32^. Since virtually all bacterial biofilms contain an EPS matrix made up of filaments that self-aggregate ^1,33–35^, similar approaches could be used for disrupting biofilms in other pathogenic bacteria. More generally, this work illustrates how targeting a biophysical mechanism, rather than a defined biochemical target, is a promising strategy that may also be targeted broadly in other infection and disease-related contexts.

## Results

### Theoretical modelling predicts small binders disrupt phage self-association

Classically biologic or synthetic binders have been used to target specific biochemical interactions between proteins and their ligands such as bacterial adhesins ^36–38^ or in targeted cancer therapies ^39^. In the case of Pf4 liquid crystal-mediated antibiotic tolerance, however, there is no specific biochemical interaction to target as liquid crystalline droplets self-assemble through depletion attraction between Pf4 filaments in crowded environments ^10,24^. In such environments, the characteristics of Pf4 droplets are governed by biophysical properties such as the size and shape of depletants (crowding biopolymers rich in the matrix) and Pf4 filaments ^30^.

We therefore used coarse-grained molecular dynamics simulations to explore how the depletion interaction between Pf4 filaments could be suppressed in crowded environments. Based on previous work on surface asperities ^40^, we hypothesised that modifying phage shape with molecules that bind to the phage filament surface could be a promising approach. We developed a minimal model and performed simulations with a mixture of phages, depletant biopolymers, and small Pf4 binders (Fig. 1 and Methods). In our simulations, Pf4 phages are modelled as hard rods (via a linear array of hard discs), depletant biopolymers are modelled as hard discs, and Pf4 binders are modelled as hard discs with a single (fixed) circular patch that is attracted to cohesive patches on the Pf4 rod (Fig. 1a). In the absence of binders, phage-phage proximity increases as depletant concentration increases (Fig. 1b), indicating that our simulations capture the depletion attraction between phages in crowded environments, consistent with published work on several filamentous phages ^10,24,30^.

**Figure 1.**
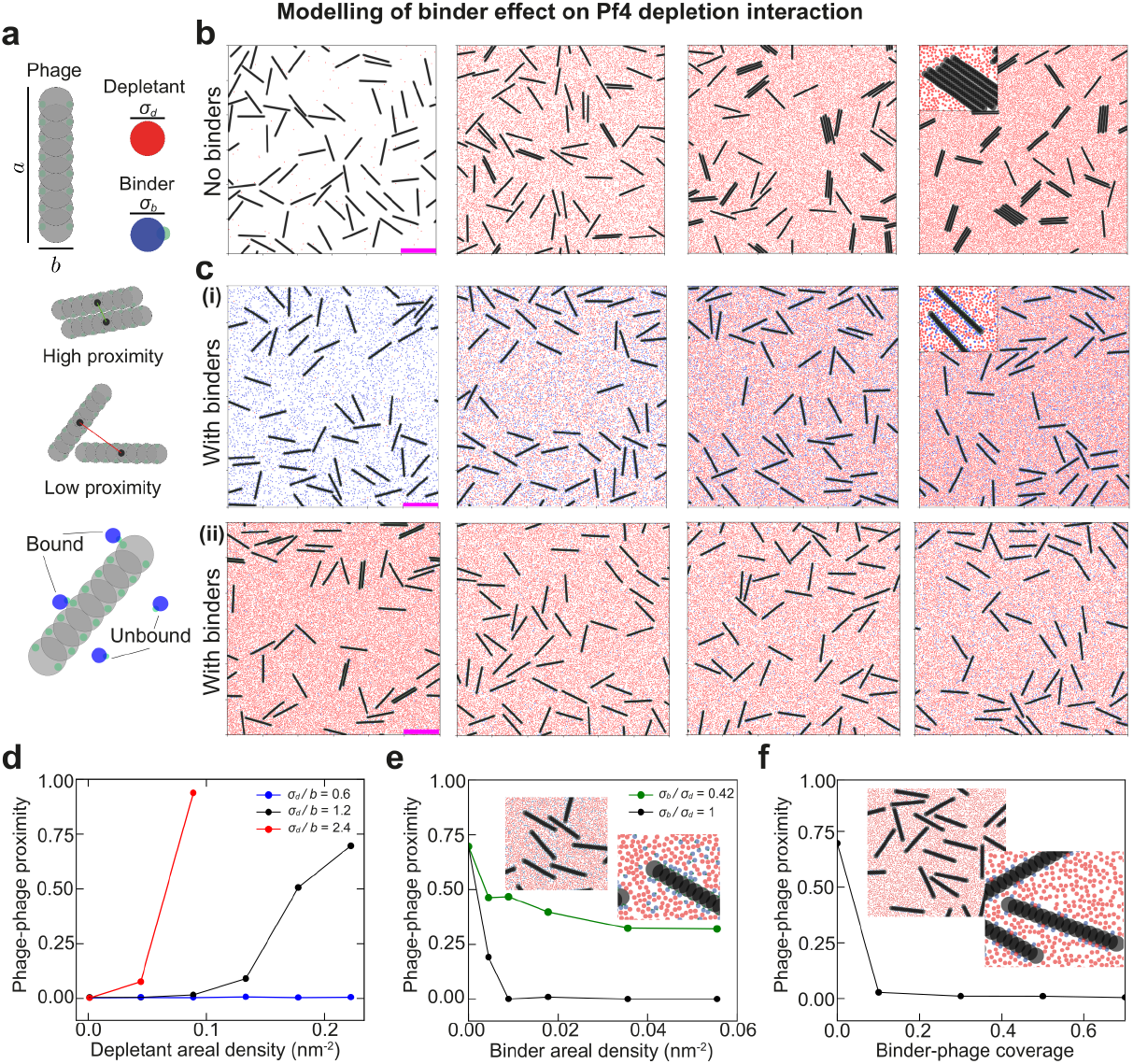
Minimal biophysical modelling suggests that Pf4 binders can disrupt depletion attraction between Pf4 phages. (**a**) Components of the coarse-grained molecular dynamics model (top): phages are modelled as hard rods of length a = 80.0 nm, and width b = 6.0 nm, the depletant particles are modelled as hard discs with a diameter σ_d_ = 2.4 nm and the binders are modelled as hard discs of diameter σ_b_ = 2.4 nm with a single (fixed) circular binding patch that is only attracted to the circular binding patches on the phage rod. Visual depiction of phage-phage alignment and localization, here termed proximity (quantified below). (**b**) Simulation snapshots of 65 phages with increasing numbers of depletant particles (100, 8000, 16000, 20000) from left to right, phages show greater proximity for increased numbers of depletant particles. Scale bar is 100 nm. (**c**) Presence of binders significantly disrupts depletion attraction between rods. (i) Same as (b) but with a constant presence of 5000 binders. Despite the increase in depletant numbers, phage proximity remains low. (ii) Increasing the numbers of binders (400, 625, 1250, and 2500) from left to right with 20000 depletant particles. Even a small number of binders disrupts the depletion interaction between rods. (**d**) Quantification of phage-phage proximity, the number of aligned phage pairs whose centers are within an approximate phage width divided by twice the number of phages, as a function of depletant density. Lines show different ratios of the depletant diameter and phage width. (**e**) Phage-phage proximity as a function of binder density for two ratios of the binder diameter and depletant diameter. Inset shows zoomed in snapshots of a simulation of 50 phages, 5000 binders, and 20000 depletant particles where the phages barely align. (**f**) Phage-phage proximity as a function of binder-phage coverage for simulations where a chosen fraction of the phage patches are permanently bound to binders. Inset shows zoomed in snapshots of a simulation of 50 phages, binder-phage coverage of 0.5, and 20000 depletant particles. In panels d-f, error bars on plots from averaging over multiple time points and simulations are too small to be visible.

By adding Pf4 binders to the simulations, we were able to suppress this depletion attraction: in the presence of the binders, phage-phage proximity is reduced in comparison to simulations without binders, across a range of depletant concentrations (Fig. 1c). Closer inspection of the rods in our simulations showed patchy and random coating of the binder along the sides of the phages (Fig. 1c). We inferred that this roughened surface reduces the overlap excluded volume between the phages, ultimately leading to a reduction in the entropic gains of the excluded depletants away from the phage surface. Our simulations suggest that for the suppression of depletion, the binder must be large enough in comparison to the depletant particles (Fig. 1d-e), in agreement with previous works on surface asperities ^40–45^. Overall, our simulations suggest that a binder could suppress the depletion attraction between phages by increasing the average roughness of phage surfaces (Fig. 1f), providing a potential mechanism to inhibit phage liquid crystalline droplets (Methods). We therefore predicted that a sub-stoichiometric Pf4 binder might be able to efficiently prevent Pf4 droplet formation, and thereby subvert the antibiotic tolerance conferred by the *P. aeruginosa* biofilm matrix.

### Development of single-domain antibodies against Pf4

Since single-domain antibodies (nanobodies) have already shown promise in disrupting *P. aeruginosa* biofilms ^37^, we followed a similar approach to generate binders against Pf4. To this end, Pf4 phage was purified from *P. aeruginosa* PAO1 biofilms and used as an antigen to inoculate alpacas for nanobody generation. Nanobodies binding Pf4 were enriched by panning against biotinylated Pf4 immobilized on beads leading to 13 positive clones. To screen for Pf4 binders, nanobodies were purified, incubated with Pf4 filaments and subjected to a co-sedimentation assay. All nanobodies co-sedimented with Pf4 filaments to varying extents confirming binding (Fig. 2a and Extended Data Figure 1a). Based on co-sedimentation behavior and sequence diversity, two nanobodies (Nb43 and Nb-D11) were carried forward for further characterization and development. Surface plasmon resonance (SPR) analysis confirmed that both Nb43 and Nb-D11 bind to Pf4, however with markedly different affinities (K_d_ = 69 ± 16 nM for Nb43 and 690 ± 60 nM for Nb-D11, Fig. 2a and Extended Data Figure 1b-e). Next, we visualized Nb43 and Nb-D11 binding to Pf4 filaments using electron cryomicroscopy (cryo-EM). Pf4 mixed with Nb43 showed Pf4 filaments with a rugged surface, with speckled patchy nanobody decoration along the length of the filament as opposed to naked filaments seen in micrographs of Pf4 alone (Fig. 2c-d). Two-dimensional (2D) class averages of Pf4 with Nb43 showed extra globular densities decorating the surface of the Pf4 filament, absent in class averages from a sample without nanobodies (Fig. 2c-d and Extended Data Figure 2a). Since the major coat protein of Pf4, CoaB, is the sole protein arranged along the entire outermost length of the phage, our analysis strongly suggests that Nb43 binds to CoaB. Interestingly, increasing Nb43 concentration resulted in Nb43 clustering on Pf4 filaments, leading to heavy decoration (Fig. 2e-f). Using the previously estimated helical symmetry of Pf4 as a starting point ^10^, we performed threedimensional (3D) reconstruction of Pf4 in complex with Nb43 (Fig. 2g-h and Extended Data Figure 2a-b). Although we could only achieve a modest resolution even with heavily decorated filaments, it allowed us to model the spatial distribution of Nb43 along the Pf4 filament. This modeling from our structural data, confirmed that the extra density, not accounted for by the Pf4 phage itself, was positioned outside the phage diameter, confirming that Nb43 binds directly to CoaB (Fig. 2g-h). In contrast to Nb43, Pf4 mixed with Nb-D11 showed no apparent nanobody decoration along Pf4 filaments in raw micrographs or in 2D class averages, suggesting either an alternative binding site for Nb-D11, or poorer binding due to lower affinity (Extended Data Figure 1d-e and 2c).

**Figure 2.**
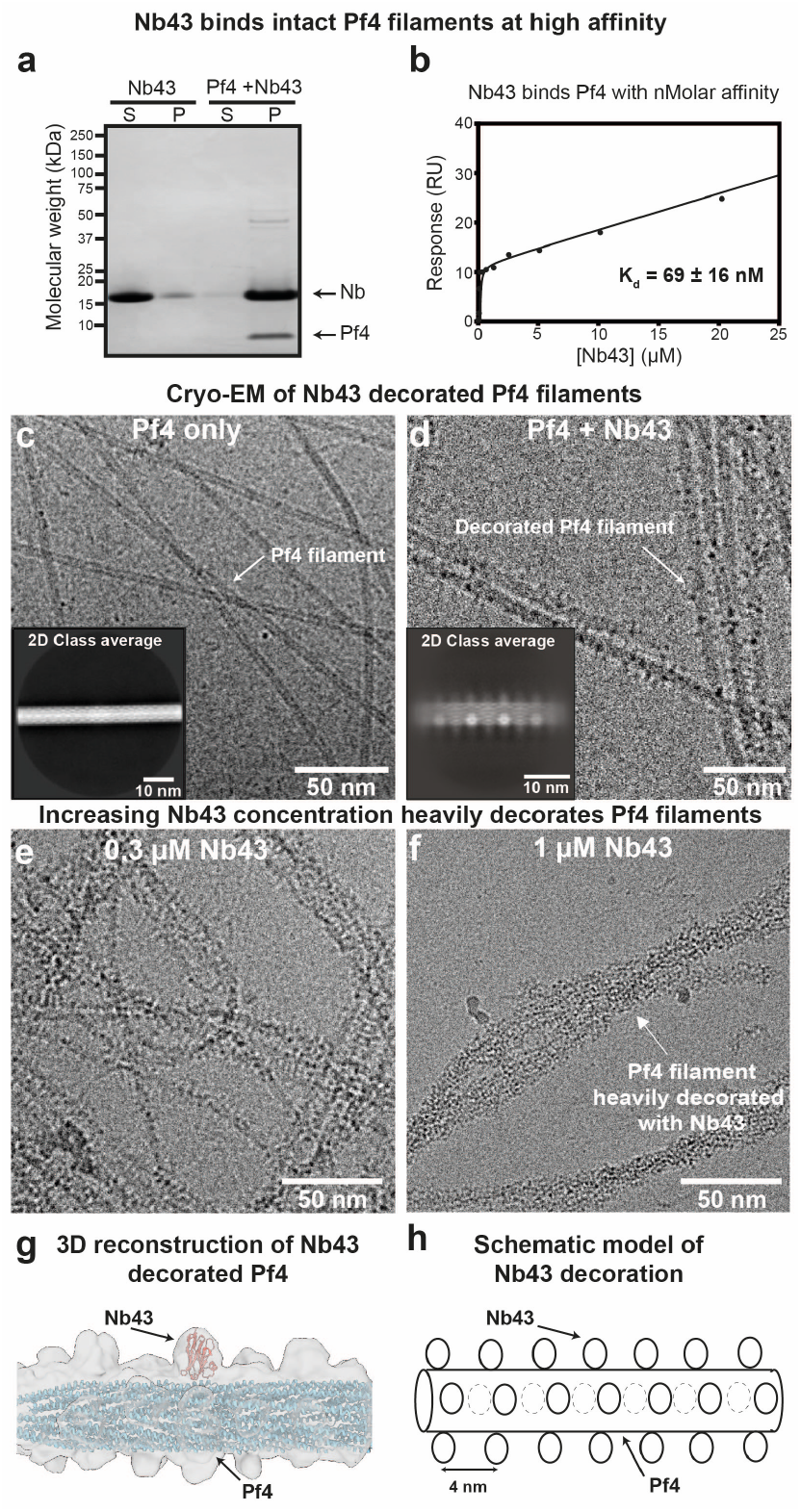
Development of a Pf4 binder along the length of the phage. (**a**) Coomassie-stained SDS-PAGE of the soluble (S) and pellet (P) fractions from Pf4 filament co-sedimentation assay with Nb43. Nb43 (1 *μ*g) was incubated alone or in the presence of Pf4 (1 *μ*g) as indicated prior to centrifugation at 100,000 g. Without Pf4 filaments, Nb43 remained predominantly in the soluble fraction, but Nb43 incubated with Pf4 co-sedimented with Pf4 in the pellet fraction after centrifugation, indicating Nb43 binding to Pf4 filaments. Arrows indicate bands at the correct molecular weight for Nb43 and Pf4 major coat protein (CoaB). Molecular weight markers are indicated on the left. (**b**) SPR response curve showing Nb43 has a high affinity for Pf4 (K_d_ = 69 ± 16 nM) (**c**) Cryo-EM micrograph of Pf4 alone, showing undecorated filaments with a diameter of ~6 nm. Inset shows representative 2D class average. (**d**) Cryo-EM micrograph of Pf4 with 0.1 *μ*M Nb43, showing Pf4 filaments decorated in a speckled manner by Nb43. Inset shows a representative 2D class average showing extra globular densities decorating the surface of the Pf4 filament. (**e-f**) Cryo-EM micrographs of Pf4 filaments incubated with increasing concentrations of Nb43, (**e**) 0.3 *μ*M Nb43 and (**f**) 1 *μ*M Nb43. Increasing Nb43 concentrations leads to increased decoration of Pf4 with nanobodies. (**g**) 3D reconstruction of Pf4 filaments with Nb43. Cryo-EM density is shown in grey with the Pf4 atomic model (blue) and Nb43 structure prediction (red) fitted into the density. (**h**) Schematic representation of Nb43 spatial distribution along a Pf4 filament, derived from (g).

### Nanobody (Pf4) binders disrupt Pf4 liquid crystalline droplet formation

Pf4 phage liquid crystalline droplets that are abundant in the biofilm EPS matrix ^9^ can be recapitulated *in vitro* by mixing purified Pf4 with biopolymers, such as alginate, a polysaccharide which is also abundant in the *P. aeruginosa* biofilm matrix ^10^. We used Alexa-488 (A488)-labelled Pf4, which allowed us to visualize the liquid crystalline droplets using fluorescence microscopy, enabling us to test the efficacy of Nb43 and Nb-D11 in preventing liquid crystalline droplet formation. We found that Nb43 was a potent inhibitor of liquid crystalline droplet formation in a dose-dependent manner, with spindle-shaped droplets almost entirely abolished at 1 μM Nb43 concentrations (Fig. 3a-c). The treated specimen showed mainly the formation of amorphous aggregates that did not resemble tactoids (Fig. 3b). Nb-D11 also had an inhibitory effect, however, a ten-fold higher concentration (10 μM) was required for a comparable decrease in droplet formation (Extended Data Figure 3a-c). Next, we assayed the ability of nanobodies to disrupt preformed droplets by premixing Pf4 phage and alginate, allowing liquid crystalline droplets to form, before addition of nanobodies. Both Nb43 and Nb-D11 could disrupt preformed liquid crystals effectively at similar concentrations to that required for prevention (1 μM for Nb43 and 10 μM for Nb-D11) (Fig. 3d-f and Extended Data Figure 3d-f). This inhibitory effect was specific against Pf4 droplets as both Nb43 and Nb-D11 did not prevent the formation of fd liquid crystalline droplets, a related filamentous phage from the Inovirus family that infects *Escherichia coli* ^30^ (Extended Data Figure 4a-f). Furthermore, incubation of Pf4 droplets with an off-target nanobody, targeting the *P. aeruginosa* CdrA adhesin ^37^, did not affect Pf4 droplet formation, showing that inhibition requires Pf4 binding, and is not due to protein crowding effects (Extended Data Figure 4g-i). Finally, both Nb43 and Nb-D11 could also disrupt Pf4 liquid crystalline droplet formation when the depletant biopolymer was changed from alginate for dextran, a biopolymer with a different charge density ^46^, further suggesting that nanobodies act through binding to Pf4 filaments without requiring a charge-based interaction with depletant molecules (Extended Data Figure 5, Methods).

**Figure 3.**
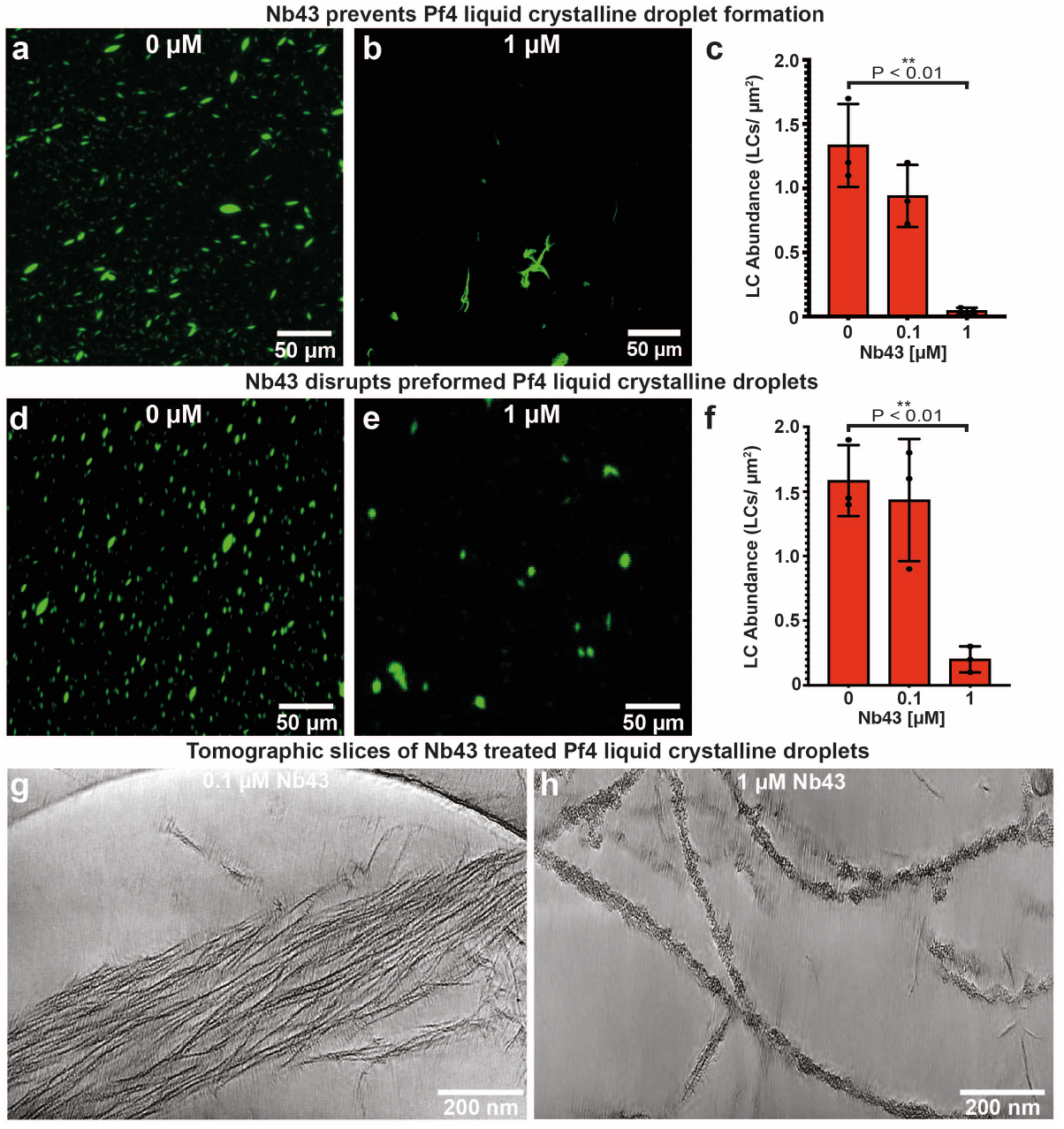
Nanobody binders are potent inhibitors of Pf4 liquid crystalline droplet formation and disrupt preformed droplets. (**a-b**) Representative light microscopy image of A488-labelled Pf4 liquid crystalline droplets formed in the presence of (a) no Nb43 or (b) 1 *μ*M Nb43. (**c**) Bar chart shows the abundance of Pf4 liquid crystals per *μ*m^2^. Addition of 1 *μ*M Nb43 results in a statistically significant reduction in droplet formation (P_value_ < 0.01). (**d-e**) Representative light microscopy images of preformed droplets treated with (d) no Nb43 (untreated) or (e) 1 *μ*M Nb43. (f) Bar chart shows the abundance of Pf4 liquid crystalline droplets per μm2 after treatment. Addition of 1 *μ*M Nb43 results in a statistically significant reduction in droplet abundance (P_value_ < 0.01). All values are derived of 30 images each from three independent biological replicates. Error bars represent standard deviation. P-values were calculated using an unpaired t-test. All images have been background subtracted. (**g-h**) Cryo-ET of Pf4 liquid crystalline droplets incubated with Nb43. Tomographic slice of a Pf4 liquid crystalline droplet specimen incubated with (g) 0.1 *μ*M and (h) 1 *μ*M Nb43.

Next, using Alexa-568 (A568)-labelled Nb43, we probed the localization of Nb43 in Pf4 droplets at low (non-inhibitory) nanobody concentrations. Nb43 was homogeneously distributed throughout the liquid crystalline droplet (Extended Data Figure 6a-d) correlating with the binding seen along single Pf4 filaments by cryo-EM (Fig. 2). Electron cryotomography (cryo-ET) revealed the ultrastructure of Nb43 incubated Pf4 droplet samples, showing that Pf4 filaments were still associated with each other at 0.1 μM Nb43 even though Nb43 decoration was visible on Pf4 filaments (Fig. 3g). This showed that Nb43 can penetrate inside the liquid crystalline droplets and bind to phages. In contrast, at 1 μM Nb43 liquid crystalline droplet formation was prevented and Pf4 filaments that were heavily decorated with clustered nanobody apparent (Fig. 3h). Parallel experiments with the other nanobody (A568-labelled Nb-D11) showed a distinct localization of Nb-D11, with Nb-D11 accumulating at the tips and edges of the spindle-shaped liquid crystalline droplets (Extended Data Figure 6e-h). In line with these observations, cryo-ET showed droplets with a nearly normal appearance with disrupted ends (Extended Data Figure 6i), confirming that Nb43 is more potent than Nb-D11, because it acts directly by binding along the phage at high affinity.

### Nanobody binders disrupt Pf4 liquid crystalline droplet-mediated antibiotic tolerance by removing antibiotic diffusion block

We next studied the effect of Nb43 on Pf4 liquid crystalline droplet-mediated antibiotic tolerance. We have previously shown addition of Pf4 liquid crystalline droplets (i.e. Pf4 with alginate) was sufficient to protect *P. aeruginosa* cells from antibiotic killing *in vitro* ^10^. As nanobody binders could prevent and disrupt liquid crystalline droplets, we hypothesized that these nanobody binders could also affect antibiotic tolerance, as Pf4 liquid crystalline droplets were previously found to be protective against antibiotic treatment ^9,10^. We tested this hypothesis by growing *P. aeruginosa* PAO1 Δ*PA0728* (lacking the Pf4 integrase) in the presence of Pf4 liquid crystalline droplets, nanobody and antibiotic. We confirmed that Pf4 droplets protect *P. aeruginosa* against aminoglycoside antibiotics in a bacterial survival assay (Fig. 4a-b). Addition of 1 μM Nb43 abolished this protective effect (P_value_ < 0.05) and reduced *P. aeruginosa* survival to the level seen in the condition with no Pf4, but alginate alone (Fig. 4a-b). Nb-D11 had a similar effect, but only at a ten-fold higher concentration (Extended Data Figure 7a-b). In control experiments, both Nb43 and Nb-D11 had no effect on bacterial growth in isolation (Fig. 4a-b and Extended Data Figure 7a-Given its higher affinity and potency against Pf4 liquid crystalline droplets, together with a defined site of binding on CoaB (Fig. 2) that mirrors our biophysical modeling (Fig. 1), and with a stronger effect in abolishing antibiotic tolerance, we went ahead with Nb43 for further experimentation.

**Figure 4.**
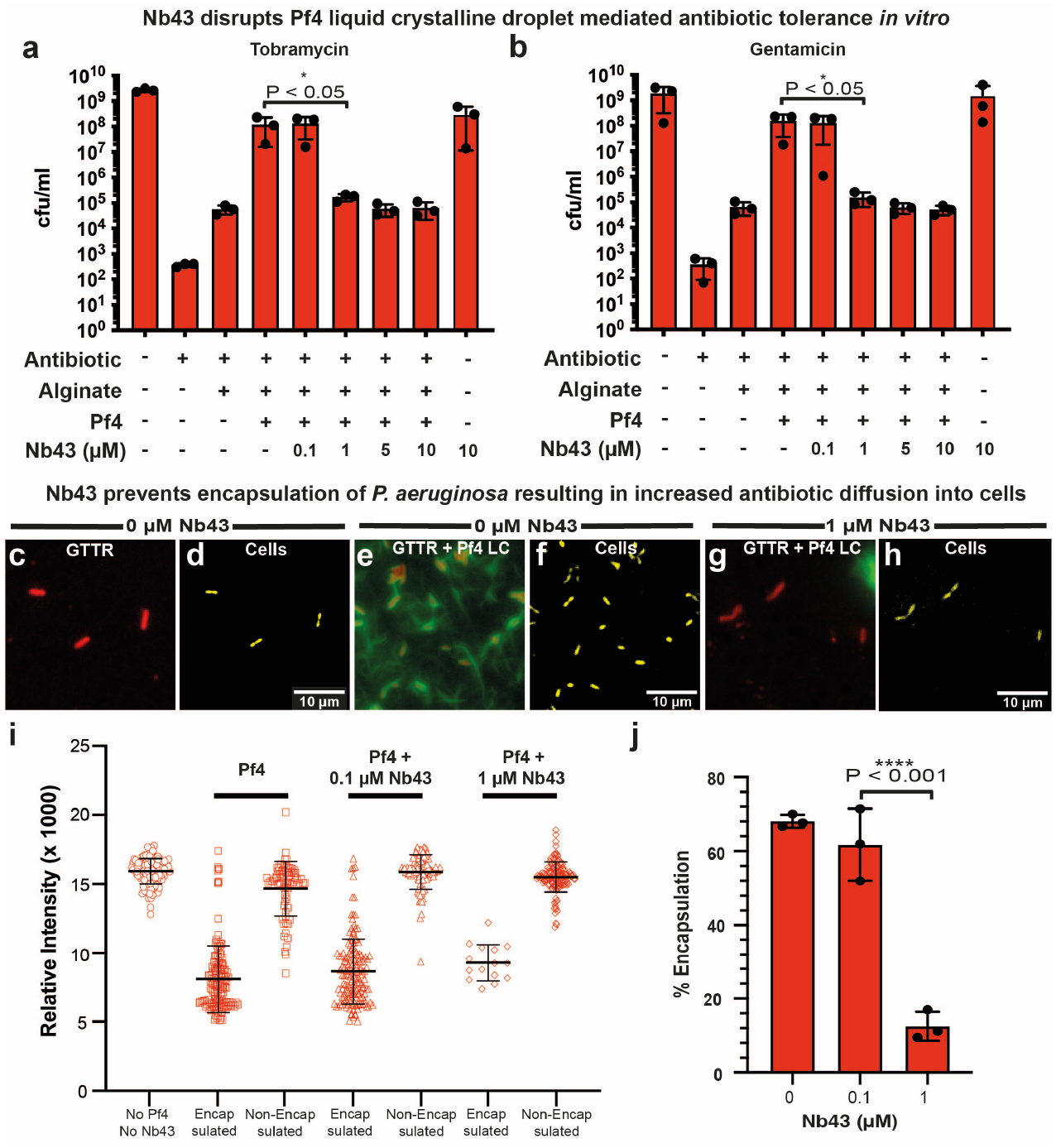
Nb43 disrupts Pf4-mediated antibiotic tolerance of *P. aeruginosa in vitro*. (**a-b**) Bar graph shows colony-forming units (cfu) per ml (y axis), a measure of *P. aeruginosa* culture cell viability after (a) tobramycin and (b) gentamicin treatment in the presence of different reagents (x axis). For both antibiotics, 1 *μ*M Nb43 significantly reduced antibiotic tolerance levels to that seen in the alginate alone condition (P_value_ < 0.05). Values shown are the mean of three independent experiments. Error bars represent standard deviation. P-values were calculated using an unpaired t-test. (**c-j**) Fluorescence and light microscopy of A488-labelled Pf4 liquid crystals incubated with or without Nb43, with Texas Red-labelled gentamicin (GTTR) incubation for 4 hours. Shown are (c, e and g) A488-Pf4 liquid crystalline droplet signal (green) and GTTR signal representing uptake of fluorescently labelled antibiotic (red) as well as (d, f and h) pseudo-colored cells (yellow) from brightfield imaging. (c-d) corresponds to a reaction without Pf4 or Nb43, (e-f) corresponds to a reaction with Pf4 but without Nb43 and (g-h) corresponds to a reaction with both Pf4 and 1 *μ*M Nb43. All images have been background subtracted. (**i**) Graph quantifying GTTR uptake in *P. aeruginosa* cells that are either encapsulated or not encapsulated by Pf4 liquid crystalline droplets after 4 hours in the presence of indicated Nb43 concentrations. Nb43 treatment resulted in increased antibiotic uptake into bacterial cells as compared to untreated specimens. (**j**) Graph plotting percentage encapsulation of bacterial cells by Pf4 liquid crystalline droplets in (i). 1 *μ*M Nb43 significantly reduces Pf4 encapsulation of bacterial cells (P_value_ < 0.001). Values shown are the mean of three independent experiments. Error bars represent standard deviation. P-values were calculated using an unpaired t-test.

Previously, we have shown that encapsulation of cells by Pf4 liquid crystalline droplets leads to a diffusion block that prevents antibiotic from reaching bacterial cells at an effective concentration ^30^. We tested whether treatment with Nb43 relieves this diffusion block: fluorescently-labelled antibiotic (Texas-Red Gentamicin – GTTR) was added to *P. aeruginosa* cells in the presence or absence of Pf4 droplets and Nb43. Antibiotic uptake was monitored using the GTTR fluorescent signal, which slowly accumulated in the cells. In control cells lacking Pf4 liquid crystalline droplets, significant antibiotic uptake into cells could be detected (Fig. 4c-d), which was diminished in the presence of Pf4 droplets, particularly in cells encapsulated by Pf4 liquid crystalline droplets (Fig. 4e-f). Addition of 1 μM Nb43 restored antibiotic uptake to levels comparable to the control with no liquid crystalline droplets (Fig. 4g-h), with a concomitant reduction in the number of cells encapsulated by Pf4 liquid crystalline droplets (Fig. 4c-j).

### Nanobody binders increase antibiotic susceptibility of *P. aeruginosa* biofilms

Given that Nb43 could affect Pf4 liquid crystalline droplet formation, antibiotic tolerance and antibiotic diffusion *in vitro*, we next tested whether this nanobody could affect antibiotic susceptibility of *P. aeruginosa* biofilms. To this end, we tested the efficacy of Nb43 in static biofilms, where biofilm growth was measured using crystal violet staining in the presence of tobramycin and varying Nb43 concentrations (Fig. 5a-b). Nb43 was added either at the beginning of growth (pre-incubation) or after 24 hours growth (without preincubation), with tobramycin being added at the 24 hours timepoint in both cases. Under pre-incubation conditions, 1 μM Nb43 significantly reduced biofilm growth by 50% compared to an antibiotic-only control at 1 μM Nb43 (P_value_ < 0.01). Without pre-incubation, a higher concentration of 2.5 μM Nb43 was required to obtain a similar reduction in biofilm growth as compared to the antibiotic treatment alone (P_value_ < 0.01) (Fig. 5a-b).

**Figure 5.**
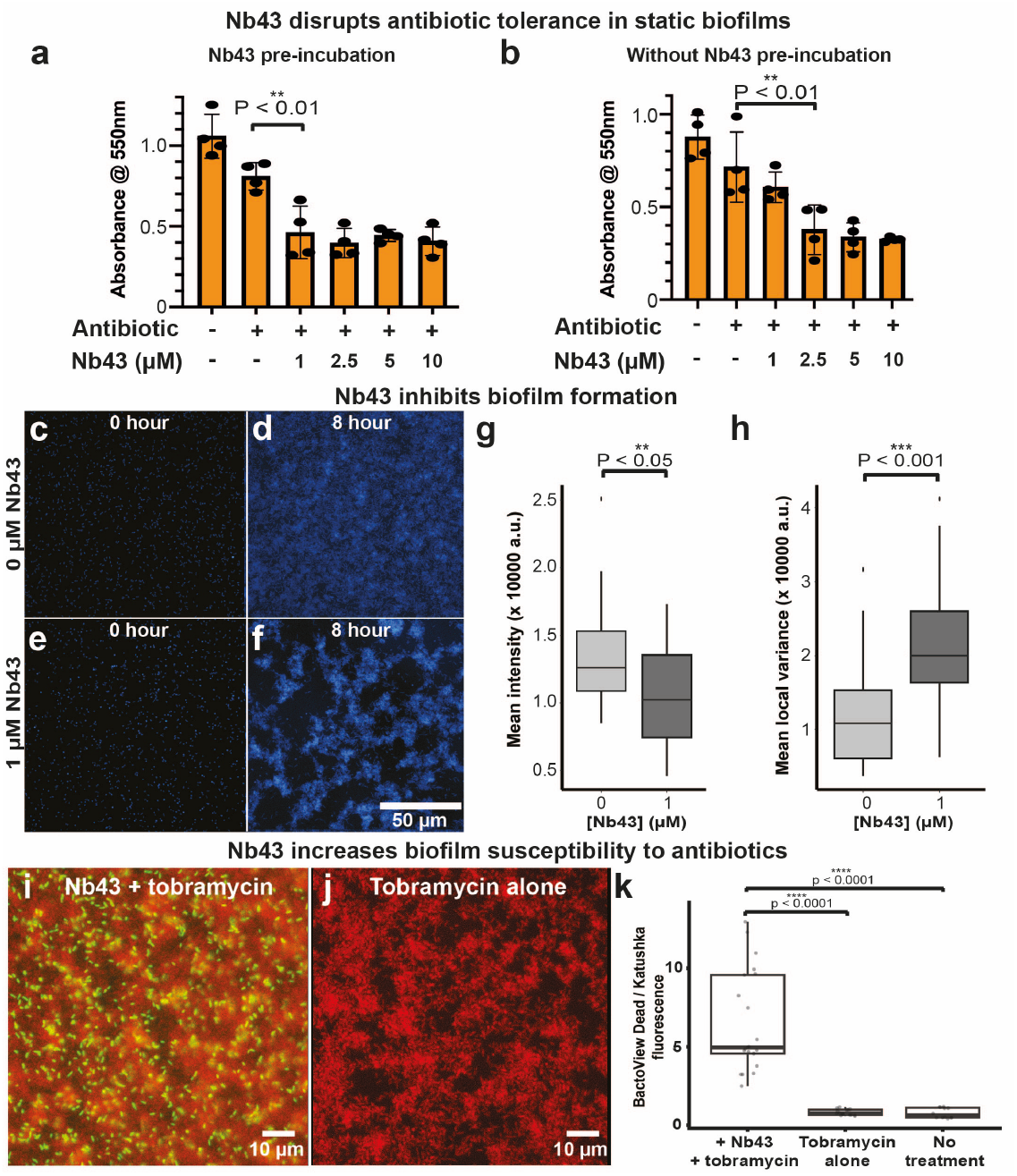
Nb43 treatment makes biofilms more susceptible to antibiotics. (**a-b**) *P. aeruginosa* PAO1 static biofilms were grown in 96-well plates and treated with Nb43 either (a) at the inoculation stage (pre-incubation) or (b) after 24 hours growth (without pre-incubation) with the indicated concentrations of Nb43. At the 24 hours timepoint, 1 *μ*g/ml tobramycin was added and cultures incubated for a further 8 hours, before plates were treated with crystal violet to assay for biofilm growth. Graphs show absorbance at 550 nm (crystal violet signal indicating amount of biofilm) on the y axis and components added to the assay on the x axis. Mean values from four replicates are plotted. Significantly less biofilm is present with (a) 1 *μ*M Nb43 in preincubation conditions (P_value_ < 0.01) and (b) 2.5 *μ*M Nb43 in the without pre-incubation condition (P_value_ < 0.01) compared to the control without Nb43. Error bars represent standard deviation. P-values were calculated using an unpaired t-test. (**c-h**) *P. aeruginosa* (PA14:mTFP, blue) biofilms cultured in a custom microfluidic device in KA biofilm medium with or without Nb43 in the culture medium. (c-f) Micrographs of mTFP intensity at the glass-liquid interface during PA14:mTFP culture over 16 hours, either in the absence of Nb43 (c-d) or in the presence of 1 *μ*M Nb43 (e-f). (g) mTFP mean intensity in images collected at the 15.5-16 hour time point. (h) Local variance of mTFP intensity across a radius of 8.1 *μ*m (50-pixel) diameter regions of images shown in (d and f), used to assay biofilm “patchiness”. The 0 *μ*M nanobody condition represents 14, and the 1 *μ*M condition represents 6 biological replicates. P-values are determined using a linear mixed effects (lmer) model that incorporates experiment date and flow cell as random effects, and Nb43 presence or absence as a fixed effect. (**i-k**) PA14 att7::Katushka2S biofilms (red) grown in flow cells with KA media for 20 hours, followed by treatment with tobramycin with or without Nb43 for 10 hours (Methods). BactoView Dead stain (green) was included to stain dead cells. (i) Biofilms (red) exposed to 2.5 *μ*M Nb43 and 10 *μ*g/ml tobramycin. (j) Biofilms exposed to tobramycin alone. (k) Graph showing the ratio of the field of view means for BactoView Dead stain (green) and Katushka2S (red) after 10 hours of treatment. The mean ratios were 6.48 ± 0.47, 0.81 ± 0.56, and 0.77 ± 0.62 for Nb43 with tobramycin, tobramycin only and untreated experiments respectively. Mean ratios were compared using a one-way ANOVA with Tukey’s post-hoc test. Three or more biological replicates were conducted for each condition and 1-2 technical replicates (flow cells) were imaged per biological replicate. Three fields of view were assessed for each technical replicate.

To explore the effect of Nb43 on biofilms further, we used a custom microfluidics flow system to cultivate *P. aeruginosa* biofilms under flow, which we have used previously to assay antibiotic treatment of *P. aeruginosa* biofilms ^37^. First, utilizing a *P. aeruginosa* strain expressing a fluorescent protein (mTFP – blue) we monitored Nb43 effect on biofilms in the absence of antibiotic. Nb43 treatment led to destabilization of biofilm growth and morphology even in the absence of antibiotic, with treated biofilms characterized by significantly less biofilm growth (P_value_ < 0.05) and a patchier cell distribution (P_value_ < 0.001) than biofilms grown in the absence of Nb43 (Fig. 5c-h). Next, we monitored the effect of Nb43 treatment on the antibiotic susceptibility of *P. aeruginosa* biofilms under flow. We used a sublethal tobramycin concentration for our experiments and monitored biofilm biomass and antibiotic susceptibility using *P. aeruginosa* cells constitutively expressing the Katushka2S fluorescent protein (far-red), together with BactoView dead staining (green) to identify dead cells. Biofilms treated with Nb43 and tobramycin showed significantly higher (P_value_ < 0.0001) cell death than biofilms treated with tobramycin alone or with no treatment (Fig. 5i-k), demonstrating that Nb43 treatment reduces antibiotic tolerance of *P. aeruginosa* biofilms. These results, taken together, confirm the efficacy of Nb43 in destabilizing *P. aeruginosa* biofilm morphology and increasing antibiotic susceptibility under various conditions (Fig. 5), in line with our original theoretical modelling prediction that a phage binder could reduce phage self-association into liquid crystalline droplets (Fig. 1).

## Discussion

Bacterial infections often proceed with biofilm formation ^18^, which is associated with an over thousandfold increase in antibiotic tolerance ^6^, thus, alternative strategies are urgently needed to disrupt bacterial biofilms. *P. aeruginosa* is a particularly problematic pathogen that is responsible for significant human morbidity and mortality in hospital settings ^13^. Previous studies have shown that nanobodies could potentiate antibiotic-killing of *P. aeruginosa* ^37^ and this study reports different nanobodies targeting an important EPS matrix molecule, Pf4, that could be used in the future development of treatments against *P. aeruginosa* biofilm infections. Pf4 assumes multiple roles during infection, including biofilm matrix stabilization ^9^, acting as an immune-decoy ^26^ and increasing antibiotic tolerance ^10^, hence nanobodies targeting Pf4 could have multiple use cases against *P. aeruginosa.* Directed evolution to increase nanobody affinity ^47^ as well as conjugation to molecules that degrade other matrix components or activate the host immune system could increase nanobody potency in the future ^48^. Furthermore, since bispecific antibodies targeting the *P. aeruginosa* biofilm matrix component Psl and Type-III secretion system protein PcrV have been tested in the clinic ^49^, similar multi-specific antibodies could be developed in the future for clinical applications.

Many bacteria including *Neisseria meningitidis* and *Vibrio cholerae* use filamentous phages from the Inovirus genus for increasing their virulence during infection ^31,32,50^, therefore similar strategies may be pursued for infection prevention in such pathogenic bacteria. Almost all bacterial biofilms contain a crowded EPS matrix made up of filaments that self-aggregate ^1,4,33–35^, thus similar approaches of disrupting the EPS matrix could be used for biofilm disruption in a variety of bacteria.

Rather than adopting the usual route of blocking an active site of an enzyme or blocking a proteinprotein interaction, in this proof-of-principle study, we have targeted a pervasive biophysical mechanism operating in biofilms, namely the entropically-driven depletion interaction between matrix filaments in crowded environments (Extended Data Figure 8). Our study was guided by biophysical modeling (Fig. 1), which suggested that a sub-stoichiometric binder could disrupt Pf4 phage liquid crystalline droplets, through surface roughening, which would dampen the depletion attraction between phage filaments. This hypothesis was realized by nanobody development, which showed clear nanobody binding with the Pf4 coat (Fig. 2), Pf4 liquid crystalline droplet disruption (Fig. 3), removal of the Pf4-mediated antibiotic diffusion block (Fig. 4) and finally substantially reducing the antibiotic tolerance of *P. aeruginosa* in biofilms (Figs. 4–5). In summary, this work illustrates that targeting biophysical mechanisms, rather than a defined biochemical target, is a promising inhibition strategy that may also be applied in other infection and disease-related contexts (Extended Data Figure 8).

## Acknowledgements

This work was supported by the Medical Research Council, as part of United Kingdom Research and Innovation (also known as UK Research and Innovation) [Programme MC_UP_1201/31 to T.A.M.B]. For the purpose of open access, the MRC Laboratory of Molecular Biology has applied a CC BY public copyright license to any Author Accepted Manuscript version arising. T.A.M.B. would like to thank the Wellcome Trust (grant 225317/Z/22/Z) and the Lister Institute for Preventative Medicine for support. P.P. was supported by a UK Research and Innovation (UKRI) Future Leaders Fellowship (MR/V022385/1). G.A.O was supported by funding from NIH/R37-AI83256. R.J.O. was supported by grants from Rosalind Franklin Institute, funding delivery partner EPSRC, Wellcome Trust (223733/Z/21/Z), BBSRC (BB/V018523/1) grants and John Fell Fund, University of Oxford. We thank VIB Nanobody Service Facility, Belgium and Jiandong Huo for raising some of the nanobodies and providing plasmids encoding nanobodies. We would also like to thank the MRC LMB electron microscopy, light microscopy, biophysics and scientific computing facilities. P.P and L.K.D. thank the computing facilities and support at the Department of Mathematics, UCL.

## Author contributions

Conceptualization, T.A.M.B. and A.T.; methodology, A.T., M.G., L.K.D., S.P., J.B., P.M., Z.W., C.M.C., J.H., R.J.O., G.A.O, P.P., T.A.M.B.; investigation, A.T., M.G., L.K.D., S.P., J.B., P.M., Z.W., C.M.C., P.P, T.A.M.B.; visualization, A.T., M.G., L.K.D., S.P., J.B., P.M., Z.W., C.M.C., T.A.M.B.; funding acquisition, T.A.M.B., R.J,O., G.A.O., P.P.; project administration, T.A.M.B., A.T., G.A.O., P.P.; supervision, T.A.M.B., A.T., R.J.O., G.O.A, P.P.; writing – original draft, T.A.M.B., A.T., L.K.D., P.P.; writing – review & editing: T.A.M.B. and A.T. along with all authors.

## Data and Materials availability statement

Further information and requests for resources and reagents should be directed to Tanmay Bharat. The nanobodies described in this paper, and where applicable their sequences, may be made available by the MRC Laboratory of Molecular Biology upon request under the terms of an MTA with restrictions on commercial use, confidentiality and publication.

## Materials and Methods

### Biophysical methods: modelling of binder inhibition of depletion interaction

To model a system of phages, depletants, and binder particles we used coarse-grained molecular dynamics simulations. We simulated a two-dimensional system consisting of *N*_*r*_phages (with either *N*_*r*_ = 65 or *N*_*r*_ = 50 depending on the simulation) of length *a* = 80 nm and width *b* = 6.0 nm. To represent each phage we used a rigid linear array of *N*_*m*_=17 overlapping discs of diameter *b* to ensure a sufficiently smooth surface (Fig. 1a). Simulations also included *N*_*d*_ depletant discs of diameter σ_*d*_, and *N*_*b*_ binder discs of diameter σ_*b*_. To mediate the interaction between phages and binders, we included two smaller discs, or patches, of diameter σ_*p*_ fixed just beneath the surface of each disc on the phages, and one patch of diameter σ_*p*_ fixed on the circumference of each binder (Fig. 1a). In the simulations, all *N* = 3*N*_*r*_*N*_*m*_ + 2*N*_*b*_ + *N*_*d*_ particles were permitted to move in a box of fixed area *L*^2^ with periodic boundary conditions at a fixed *N* and temperature *T* = 297 K (canonical ensemble).

In our model, the non-patch particles experience excluded volume (steric) interactions, with the only attractive interaction occurring between the patch particles on each disc of the rod and the single patch on the binders. Any interaction between the patches on the rod are zero by construction. The total (sum over all pairs of particles) potential energy *V* accounting for these interactions is given as

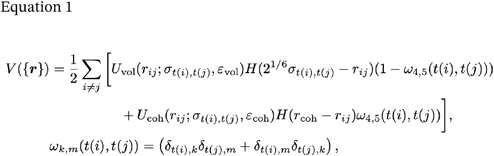

where {***r***} is the set of all positions, *r*_*ij*_ = |r_i_ − r_j_| is the distance between two particles *i* and *j*, ε_*vol*_ = 10 *k*_*B*_*T* is the excluded volume interaction strength, *ε*_*coh*_= 5 *k*_*B*_*T* is the rod-patch to binder-patch cohesion strength, *r*_*coh*_= 3σ_*p*_= 1.5 nm is the cohesion range, *t*(*i*) ∈ (1,2,3,4,5) denotes the particle type of *i* (1 = rod, 2 = depletant, 3 = binder, 4 = binder-patch, and 5 = rod-patch), factors of ω_*k,m*_(*t*(*i*), *t*(*j*)) is a way to only impose excluded volume interactions to types 1 to 3 and cohesive interactions to types 4 to 5, δ_*i,j*_ is the Kronecker delta function, *H*(…) is the Heaviside function, and σ_*t*(*i*),*t*(*j*)_ = (σ_*t*(*i*)_ + σ_*t*(*j*)_)/2 is the Lorentz-Berthelot rule with σ_*t*(*i*)_ being the diameter of a particle of type *t(i). U*_vol_ imposes excluded volume and is given by the Weeks-Chandler-Andersen potential as:

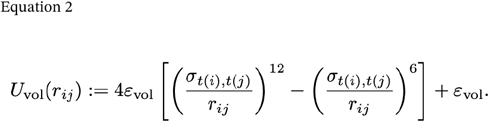

The cohesive interaction, *U*_coh_, is determined by a truncated and shifted Lennard-Jones potential given as:

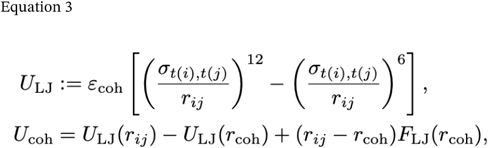

where *F*_LJ_ is the force of the Lennard-Jones potential.

To obtain dynamical trajectories of the above model, we used the following protocol. We time integrate overdamped Langevin dynamics in the LAMMPS (Aug 2024) package ^51^, with a timestep of δ*t* = 0.002 ns. We impose the rigidity of the rods through a constraint term in the dynamics, as specified by the *fiix rigid* command in LAMMPS. Initially, the system is prepared with the rods uniformly spread across the plane in grid-like fashion, and then a small initial simulation is conducted dragging depletant and binder particles into the simulation box of area *L*^2^= 600×600 nm^2^. Before any recording of the observables, the system is equilibrated for 10^7^ timesteps and production runs last 5 × 10^7^ timesteps. Equilibration is checked through inspection of the total energy and the averaged orientation of the rods.

### Biophysical modelling considerations: mechanism of binder inhibition of Pf4 liquid crystalline droplet formation

In previous work, the phage Pf4 has been found to form protective liquid-crystalline droplets around bacterial cells in environments that are charge-screened and crowded; such environments suppress charge-based phage-phage repulsion and generate a depletion attraction between phages ^9,10^. Therefore, in this study, we applied our model to explore a method to suppress a non-specific interaction induced by crowding agents, rather than adopting the usual route of blocking an active site of an enzyme or blocking a protein:protein interaction. As described in the main text, our biophysical modelling results (Fig. 1) suggested that a sub-stoichiometric binder could disrupt Pf4 phage liquid crystalline droplets through surface roughening, which dampens the depletion attraction. Our model predicts that such a mechanism would require the binder being large enough in comparison to the depletant (Fig. 1e). In our experiments we found that Nb43, a nanobody that binds to Pf4 and decorates its surface (Fig. 2), suppresses the formation of liquid crystalline droplets (Fig. 3), in line with our theoretical prediction of reduced Pf4 self-association because of surface roughening. At higher concentrations of Nb43, we observed clustering of the binder on Pf4 filaments, leading to heavy decoration (Fig. 2e-f), which may contribute to the effect of the surface roughening by increasing its effective size.

In addition to the surface roughening mechanism, the nanobody Nb43 may also reduce the effective size of the depletant, alginate, in the phage-phage depletion interaction. This would involve the phage-bound nanobodies screening charge-based repulsion between phages and depletants (both alginate and phages are negatively charged); however, because our experiments were performed in a charge-screened environment, it is unlikely that this is a major factor in the nanobody’s mechanism. Furthermore, the nanobody’s action was found to be replicated in experiments with dextran as a depletant, which carries less overall negative charge (Extended Data Figure 5). Overall, the combined modelling and experimental results suggest that suppression of depletion via surface roughening, as predicted by our model, is the main mechanism by which the nanobody Nb43, a Pf4 binder, disrupts Pf4 liquid crystalline droplet formation.

### Bacterial strains

*P. aeruginosa* strains PAO1 or PAO1 Δ*PA0728* (a kind gift from Prof. Patrick Secor, Montana State University) or PA14:mTFP (modified to express teal fluorescent protein (mTFP) under the PA1/04/03 promoter at the *att7* site) or the PA14 att7::Katushka2S were utilised as indicated. *E. coli WK6* were used for periplasmic expression of nanobodies. Shaking cultures of bacteria were grown at 37 °C in Luria-Bertani (LB) medium with agitation at 180 rpm (revolutions per minute) unless otherwise specified.

### Bacterial media

The following bacterial media was used for experiments. Luria-Bertani (LB): Composed of 5 g/L yeast extract, 10 g/L tryptone, and 5 g/L NaCl, dissolved in milliQ water. Terrific broth (TB): 12 g/L tryptone, 24 g/L yeast extract, 0.4 % (v/v) glycerol, 0.017 M potassium dihydrogen phosphate and 0.072 M potassium phosphate dibasic. KA biofilm medium ^52^: Composed of 6 g/L Tris-HCl (adjusted to pH 7.4), 4 g/L L-arginine HCl, 174 mg/L dipotassium sulfate, and 72 mg/L magnesium sulfate in water.

### Pf4 phage amplification

Pf4 bacteriophage was produced as described previously ^10^. Briefly, Pf4 was amplified from stock Pf4 isolated from PAO1 biofilms ^10^ by incubating 1 × 10^3^ pfu (plaque forming units)/ml of Pf4 stock with 1 ml of PAO1 culture at 0.5 optical density measured at 600 nm wavelength of light (OD_600_) for 15 minutes before mixing with hand-hot 0.8 % (w/v) agar. The mixture was plated onto 10 cm^2^ LB-agar plates and incubated overnight at 37 °C. Each plate was covered with 5 ml phosphate buffered saline (PBS) and incubated for 6 hours at room temperature (RT) before the PBS was collected and centrifuged (12,000 *g*, 30 minutes, 4°C) to remove debris. Pf4 was precipitated from the supernatant by polyethylene glycol (PEG) precipitation. The supernatant solution was adjusted to 0.5 M NaCl and Pf4 precipitated by addition of 10 % (w/v) PEG 6000 (Sigma) followed by overnight incubation at 4 °C. Precipitated Pf4 phage particles were pelleted by centrifugation (12,000 *g*, 30 minutes, 4 °C), resuspended in PBS and dialysed overnight against PBS using 10 kDa MWCO (molecular weight cut-off) snakeskin dialysis membranes (Thermo Fisher Scientific). Yield of the Pf4 bacteriophage preparation was estimated using Nanodrop (Thermo Fisher Scientific).

### Nanobody generation, expression and purification

Nb-D11 was generated by the Nanobody Discovery Platform, Rosalind Franklin Institute, UK and Nb32-45 were generated by the VIB Nanobody Service Facility, Belgium by methods previously published ^53,54^. Plasmids encoding His_6_-tagged-nanobodies were transformed (separately) by heat shock into *E. coli WK6* cells. Terrific broth (TB) supplemented with 100 *μ*g/ml ampicillin was inoculated with transformed *E. coli WK6* cells and incubated at 37 °C with agitation at 180 rpm. Upon reaching an OD_600_ of between 0.6-0.8, cultures were induced with 1 mM isopropyl β-D-thiogalactoside (IPTG) and incubated overnight at 23 °C with agitation at 180 rpm. Cells were pelleted by centrifugation (5000 *g*, 10 minutes, 4 °C) and resuspended in 15 ml ice-cold Tris/Ethylenediaminetetraacetic acid (EDTA)/Sucrose (TES buffer – 200 mM Tris, 650 μM EDTA and 500 mM sucrose) per litre of culture and incubated for 1 hour with agitation on an orbital rocker. To disrupt the periplasm, 30 ml ice-cold TES/4 (1 part TES buffer and 3 parts milliQ water) was added per litre of culture and agitated on an orbital rocker for 45 minutes at 4 °C. After incubation, debris was pelleted by centrifugation (10,000 *g*, 30 minutes, 4 °C), the supernatant recovered and subjected to Nickel-NTA affinity chromatography with a 5 ml His-Trap high performance (HP) column (Cytiva) followed by size exclusion chromatography with a Superdex S75 column (Cytiva) in size exclusion buffer (50 mM Tris (pH 7.4), 150 mM NaCl, 0.5 mM TCEP). Nanobody containing fractions were concentrated and the protein concentration was estimated using a nanodrop (Thermo Fisher Scientific). Nanobodies were aliquoted, snap-frozen in liquid nitrogen and stored at −80 °C ahead of use.

### Nanobody co-sedimentation with Pf4 filaments

Nanobody was centrifuged (100,000 *g*, 1 hour, 4 °C) to remove any aggregates formed during storage and the supernatant used in the co-sedimentation assay after further protein concentration determination by nanodrop (Thermo Fisher Scientific). Each nanobody (1 *μ*g) was incubated either alone or with an equal amount (w/w) of purified Pf4 in a 100 *μ*l reaction volume for 1 hour at room temperature before centrifugation (100,000 *g*, 1 hour, 4 °C). The supernatant was carefully removed from the pellet, which was resuspended in the same volume of PBS as the supernatant. Equivalent volumes of the supernatant (S) and pellet fractions were loaded onto 4-12% tris-glycine Novex gels (Thermo Fisher Scientific) and proteins visualised by Coomassie Blue staining.

### Surface Plasmon Resonance (SPR)

SPR was performed using a Biacore T200 using C1-sensor chip (Cytiva). Both reference control and analyte channels were equilibrated HBS-N buffer (Cytiva). Pf4 was immobilised onto the chip surface *via* amide coupling using the supplied kit (Cytiva) to reach a RU value of 400 RU. SPR runs were performed with analytes injected for 120 seconds followed by a 300 second dissociation in a 1:2 dilution series with initial concentrations of 10 *μ*M. After reference and buffer signal correction, sensorgram data were fitted using Prism Graphpad (GraphPad Software Inc). The equilibrium response (R_eq_) data were fitted to a single site interaction model with linear non-specific binding to determine *K*_d_:

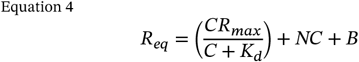

where C is the analyte concentration and R_max_ is the maximum response at saturation, N is the non-specific binding and B is the background resonance.

### Cryo-EM grid preparation

Samples for cryo-EM were prepared as described previously ^10^, by pipetting 2.5 *μ*l of the sample onto freshly glow-discharged Quantifoil grids (Cu/Rh R2/2, 200 mesh or Au R1.2/1.3, 300 mesh for cryo-EM, Cu/Rh R3.5/1, 200 mesh for cryo-ET) and plunge frozen into liquid ethane in a Vitrobot Mark IV (Thermo Fisher Scientific). Plunge-frozen grids were transferred to liquid nitrogen and stored until imaging.

### Cryo-EM and cryo-ET data collection

Cryo-EM data for screening specimens were collected either using a Titan Krios or Glacios microscope (Thermo Fisher Scientific) operated at 300 kV or 200 kV respectively. High-throughput data was collected using the EPU software on a Titan Krios microscope fitted with a Selectris energy filter (slit width 10 eV) and a Falcon 4i direct electron detector operating in counting mode at an unbinned, calibrated pixel size of 0.96 Å. A combined total dose of approximately 50 e^-^/Å^2^ was applied with each exposure lasting 4 seconds and 40 frames were recorded per movie. In total 3601 movies were collected between −1 to −2.5 *μ*m defoci. Tilt series data for cryo-ET was collected on a Titan Krios using the BioQuantum energy filter (Gatan) and K3 direct electron detector (Gatan) with the SerialEM software^55^. Tilt series were collected in a dose-symmetric scheme starting from 0° between ± 60° with 1° tilt increment between −6 to −8 *μ*m defoci, with a combined dose of approximately 163 e^-^/Å^2^ applied over the entire series. Tilt series were collected at an unbinned, calibrated pixel size of 2.13 Å. Acquired tomographic data were pre-processed in RELION5 ^56^ and tomograms were reconstructed using AreTomo ^57^ and denoised using cryoCARE ^58,59^.

### Cryo-EM data processing

Cryo-EM image processing of 2D image data of nanobody-decorated phage filaments was performed in cryoSPARC ^60^. After patch motion correction and CTF estimation, filament positions were determined within micrographs using the cryoSPARC filament picker, extracting particles every 1.2 nm along the filament axis, resulting in one unique nanobody binder in each box according to estimates based on previous 2D class averages. 2D class averages were then created in cryoSPARC. A wide range of symmetries for helical refinement was tested but did not yield densities with resolved Nb43-bound CoaB subunits. 3D reconstructions of Nb43-bound Pf4 were ultimately generated using helical refinement in cryoSPARC with no symmetry applied.

### Fluorescent labelling of Pf4 and nanobody

Purified Pf4 or nanobody was dialysed into PBS using 10 kDa MWCO snakeskin dialysis membrane (Thermo Fisher Scientific). To adjust the pH of the sample, 100 μl of 1 M sodium carbonate buffer pH 9.2 was added to 900 μl Pf4 phage (5 mg/ml) or 900 μl nanobody (10 mg/ml). The pH-adjusted Pf4 sample was incubated with 100 *μ*g Alexa fluor 488-NHS fluorescent dye (Thermo Fisher Scientific) for 1 hour at room temperature (RT) with end-over-end agitation. The pH-adjusted nanobody samples were incubated with 100 *μ*g Alexa fluor 568-NHS fluorescent dye (Thermo Fisher Scientific). Both A488-labelled phage and A568-labelled nanobody were isolated from free dye by passing over two PD10 desalting columns (Cytiva). Protein concentration was estimated by nanodrop (Thermo Fisher Scientific).

### Fluorescence microscopy

#### Pf4 liquid crystalline droplets in presence of unlabelled nanobody

A488-labelled Pf4 phage (final concentration 1 mg/ml) were mixed with sodium alginate (final concentration 4 mg/ml) or dextran (final concentration 10 mg/ml) in the presence or absence of indicated nanobody and incubated at room temperature overnight for liquid crystalline droplet prevention experiments. For disruption of preformed liquid crystalline droplets, A488-labelled Pf4 phage and alginate were incubated for 4 hours before the addition of nanobodies at the indicated concentrations and incubated at room temperature overnight. 5 *μ*l of the resulting sample was pipetted onto 1.5 % (w/v) agar pads constructed using 15 × 16 mm Gene Frames (Thermo Fisher Scientific) following the manufacturer’s protocol, with a coverslip placed on top. The slide was imaged using a Zeiss Axioimager M2 (Carl Zeiss) microscope in fluorescence mode. Quantification of individual Pf4 liquid crystalline droplet area was performed using MicrobeJ ^61^. Maxima corresponding to individual Pf4 liquid crystalline droplets were analysed using MicrobeJ shape analysis functions to quantify area. Experiments were performed in triplicate. Presented images were background-subtracted and figure panels were prepared using Fiji ^62^. Graphs were plotted using Prism Graphpad (GraphPad Software Inc).

#### Pf4 liquid crystalline droplets in presence of A568-labelled nanobody

A488-labelled Pf4 phage (final concentration 1 mg/ml) was mixed with sodium alginate (final concentration 5 mg/ml) in the presence of the indicated concentration of A568-labelled Nb43 and A568-labelled Nb-D11 and incubated at room temperature overnight. 5 *μ*l of the resulting sample were pipetted onto 1.5 % (w/v) agar pads constructed using 15 × 16 mm Gene Frames (Thermo Fisher Scientific) following the manufacturer’s protocol, with a coverslip placed on top. The slide was imaged using a Zeiss Axioimager M2 (Carl Zeiss) microscope in fluorescence mode. Presented images were background-subtracted. Fluorescence intensities of both Pf4 and nanobodies were calculated by line profile analysis by drawing a line through the liquid crystalline droplet and using the plot profile function in Fiji ^62^. Graphs were plotted using Prism Graphpad (GraphPad Software Inc).

### Texas Red-gentamicin (GTTR) diffusion into bacterial cells

PAO1 Δ*PA0728* cells were grown to an OD_600_ of 0.5 and incubated with A488-labelled phage (final concentration 1 mg/ml) and sodium alginate (final concentration 4 mg/ml) and 1 *μ*M Nb43 for 1 hour. Texas Red-gentamicin (AAT Bioquest) was added to a final concentration of 1 *μ*M and incubated for a further 4 hours. 5 *μ*l of the sample were pipetted onto 1.5 % (w/v) agar pads constructed using 15 × 16 mm Gene Frames (ThermoFisher) with a coverslip applied before subsequent imaging using a Zeiss Axioimager M2 microscope (Carl Zeiss) in both brightfield and fluorescent mode. Presented images are background-subtracted and figure panels were prepared using Fiji ^62^. Images were quantified by semi-automated segmentation of bacterial cells and associated Pf4 liquid crystalline droplet (see below) and fluorescent intensity in the Texas Red channel was measured at the coordinates of segmented bacteria. Graphs were plotted using Prism Graphpad (GraphPad Software Inc).

### Semi-automated segmentation of images with bacterial cells encapsulated by Pf4 droplets

Bacterial cells were selected from the brightfield channel images to locate the positions of cells. Bacterial cell shapes were found using the *activecontour* algorithm in MATLAB ^63^. The regions of identified bacteria were dilated and used as seed inputs for the segmentation of the Pf4 liquid crystalline droplets in the fluorescence channel using the *activecontour* algorithm. The segmented bacterial cells from the brightfield and the Pf4 liquid crystalline droplets from the fluorescence channel were then used to calculate encapsulation. Fluorescent intensity in the Texas Red channel was measured at the coordinates of segmented bacteria.

### Nanobody disruption of Pf4 liquid crystalline droplet-mediated antibiotic protection *in vitro*

An overnight culture of PAO1 Δ*PA0728* was grown in LB media at 37 °C, diluted 1 in 100 into fresh LB medium and grown at 37 °C to an OD_600_ of 0.5. 100 *μ*l of the resulting culture was added to a 96-well plate and incubated further for 30 minutes at 37 °C. 100 *μ*l of Pf4 and/or alginate, and nanobody were added to the culture such that final concentrations of components were: sodium alginate (Scientific Laboratory Supplies) (4 mg/ml), Pf4 (1 mg/ml) and nanobody at the indicated concentration. Additionally, tobramycin (10 *μ*g/ml) (Sigma) or gentamicin (10 *μ*g/ml) (Sigma) was added as indicated and cultures were grown further for 3 hours. A 10 *μ*l sample for each assay condition was serially diluted 10-fold and 100 *μ*l of the dilutions plated onto LB agar plates. Plates were inverted and incubated overnight at 37 °C and colonies forming units (cfu) enumerated. Experiments were performed in triplicate. Mean cfu/ml with standard deviation were calculated and plotted using the Prism GraphPad software (GraphPad Software Inc).

### Antibiotic susceptibility of static *P. aeruginosa* biofilms

Static biofilm assays were adapted from a previously described protocol ^64^. An overnight culture of PAO1 Δ*PA0728* was grown in LB media at 37 °C. The culture was diluted 1 in 100 into fresh LB media and 100 *μ*l pipetted into each well of a flat-bottomed 96-well plate. In the case of pre-incubation experiments, nanobody at the indicated concentration was added to the well. Plates were incubated overnight at 37 °C before addition of 0.5 *μ*g/ml tobramycin and incubation for a further 8 hours. For experiments without a pre-incubation, Nb43 was added at the indicated concentration at the same time as tobramycin. Subsequently, planktonic cells were removed from the plate by inverting and shaking out the culture. The plate was washed three times by submerging in milliQ water and shaking out contents over paper towels. To visualize biofilms, 200 *μ*l 0.1 % crystal violet (Sigma) was added to each well and incubated at room temperature for 15 minutes. Plates were washed a further three times with MilliQ water as described previously and plates were inverted and left to dry overnight. To quantify biofilm formation, 200 *μ*l 30% (v/v) acetic acid was added to each well and incubated for 15 minutes. Samples were transferred to a fresh 96 well plate and crystal violet signal was quantified by measuring absorbance at a wavelength of 550 nm in a Tecan Infinite M200 Pro plate reader. Experiments were repeated four times. Mean absorbance with standard deviation were calculated and plotted using Prism GraphPad software (GraphPad Software Inc).

### Nanobody effect on *P. aeruginosa* biofilms grown under flow conditions

#### Flow cell fiabrication

Microfluidic flow cells were fabricated using standard soft photolithography techniques ^65^, as described briefly here: To create a negative-relief mold, a silicon wafer was photolithographically patterned with SU8 3050 photoresist (Kayaku). Sylgard 184 polydimethylsiloxane (PDMS) (DOW) was cast against this mold and sectioned into individual flow cell devices. Inlet and outlet ports with a 0.75 mm diameter were punched into the PDMS before assembly. The devices were plasma bonded to 25 mm × 60 mm 1.5 cover glasses. The resulting flow cells had channel dimensions of 50 *μ*m height, 1000 *μ*m width, and 8500 *μ*m length.

#### Flow cell culture – biofiilm establishment

The method for conducting the flow cell experiments is adapted from previous work ^37,66^ and described as follows: The flow cell experiments used a *P. aeruginosa* PA14 strain modified to express teal fluorescent protein (mTFP) under the PA1/04/03 promoter at the *att7* site, this strain was used in prior flow cell experiments ^37^. Bacterial cultures were grown in LB to an OD_600_ of 2 ± 0.25, washed, and resuspended in KA medium to achieve a final OD_600_ of 0.2. These bacterial suspensions were injected into flow cells and incubated undisturbed for a 30 min attachment period. After the attachment period, syringes (1 ml) containing KA medium, with or without 1 *μ*M Nb43, were connected to the flow cell inlets via microtubing. The system was operated with a syringe pump set to 40 *μ*l/hour. Imaging was conducted using a Nikon Ti microscope equipped with a 40× objective, capturing transmitted light and CFP fluorescence channels. Images were acquired hourly over a 16-hour period, with three distinct fields of view per flow cell. Micrographs taken at 15.5– 16 hours post-initiation of flow were analyzed to evaluate the biofilm formed. Fields of view disrupted by bubbles or loss of focus were excluded. Remaining images were converted to TIFF format for analysis in Fiji ^37^. The mean intensity of each TIFF file was measured using the “Measure” tool. To quantify local variance, the “Variance” filter was applied with a radius of 8.1 *μ*m (50 pixels) diameter, generating a new image. The mean intensity of the filtered image, representing local variance, was measured with the “Measure” tool.

#### Flow cell culture – biofiilm treatment

For biofilm treatment flow cell experiments, a *P. aeruginosa* PA14 att7::Katushka2S strain was used. Katushka2S is a fluorescent protein with far-red fluorescence (588 nm Ex / 635 nm Em) ^67^. Bacterial cultures were grown in LB at 37 °C for 16 hours, before diluting 1:50 in KA biofilm medium and incubating for 4 hours at 37 °C. At 4 hours, the subculture was injected into flow cells and incubated undisturbed for a 10 minute attachment period. After the attachment period, syringes (5 ml) containing KA medium were connected to the flow cell inlets via microtubing. KA medium was introduced to the flow cells for 20 hours at a flow rate of 100 *μ*l/hour. The flow cells were not monitored via microscopy for the biofilm development period. At 20 hours the flow cells were transferred to the microscope stage and the syringes were replaced with 1 ml syringes containing KA medium with 1x BactoView™ Dead 500/515 stain (Biotium), KA medium with dead stain and 10 *μ*g/ml tobramycin, or KA medium with dead stain, tobramycin, and 2.5 *μ*M Nb43. The medium was delivered at a flow rate of 50 *μ*l/hour and the devices incubated at 37 °C. Imaging was conducted using a Nikon Ti microscope equipped with a 40X objective, capturing transmitted light, GFP fluorescence, and the Texas Red fluorescence channels. Images were acquired hourly over a 10-hour period, with three distinct fields of view per flow cell. Micrographs taken 10 hours after the initiation of tobramycin ± Nb43 treatment were analyzed to evaluate the extent of killing, indicated by BactoView™ Dead 500/515 staining. The mean fluorescence intensity for the two observed fluorescence channels, Texas Red (viable bacteria) and GFP (dead bacteria), were determined as described in the previous section. After measurement, the average GFP intensity was divided by the average Texas Red intensity for each collected field of view, to determine the ratio of dead stain signal to Katushka2S signal.

**Extended Data Figure 1.**
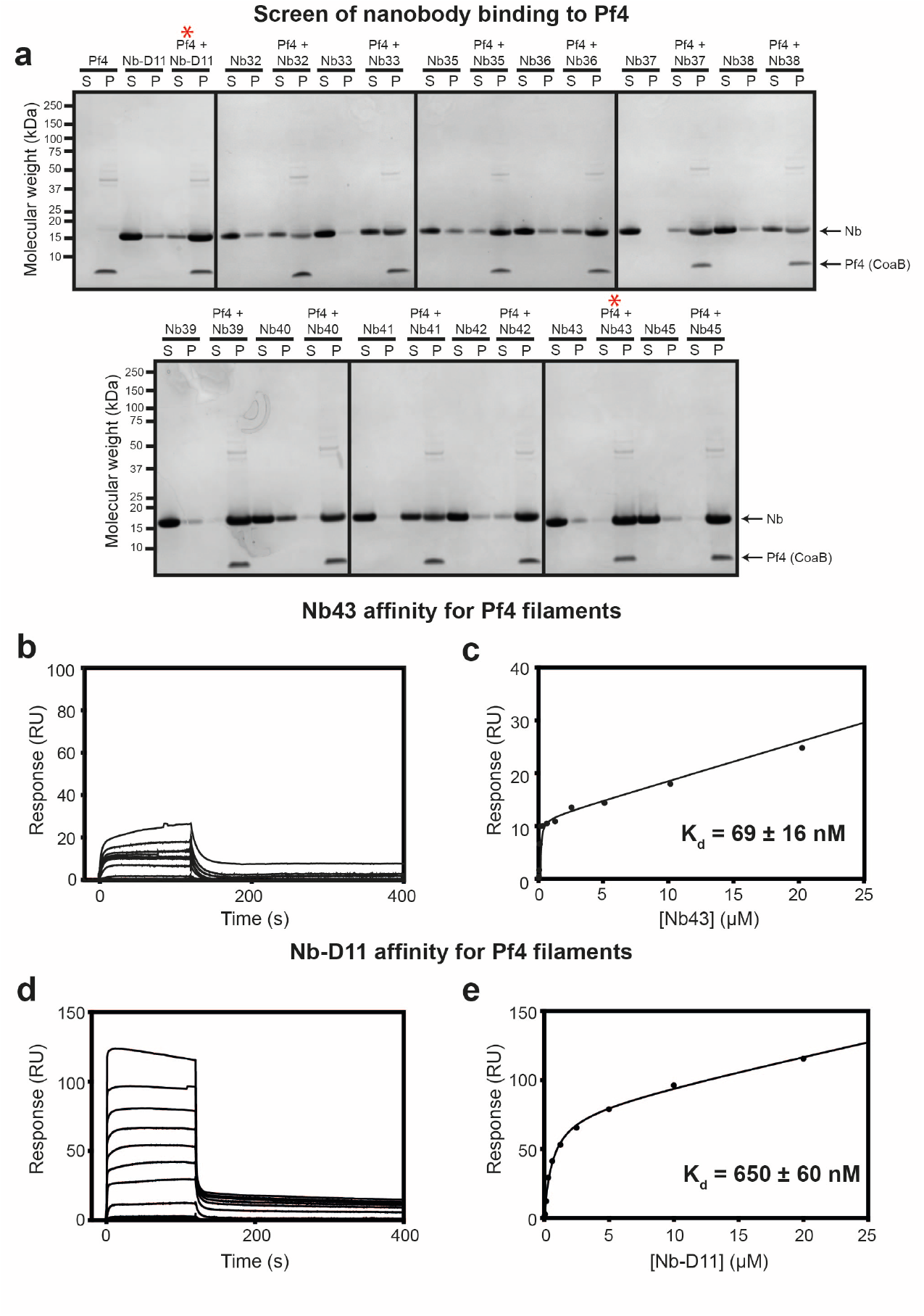
Nanobody binding to Pf4 filaments. (**a**) Coomassie-stained SDS-PAGE of the soluble (S) and pellet (P) fractions from Pf4 filament co-sedimentation assay with a panel of recombinant nanobodies. Nanobodies (1 *μ*g) were incubated alone or in the presence of Pf4 (1 *μ*g) as indicated prior to centrifugation at 100,000 g. Without Pf4, nanobodies remained predominantly in the soluble fraction, but nanobodies incubated with Pf4 co-sedimented with Pf4 in the pellet fraction after centrifugation, indicating binding to Pf4. Arrows indicate bands corresponding to nanobody (Nb) and the Pf4 major coat protein (CoaB). Molecular weight markers are shown on the left. Red asterisks denote nanobodies selected for further characterization (Nb43 and Nb-D11). (**b-e**) Pf4 was immobilized via amide coupling onto a C1-sensor chip and nanobodies were flowed over the chip. (b) Nb43 sensorgram and (c) response curve showing Nb43 has a low nanomolar affinity for Pf4 (K_d_ = 69 ± 16 nM), (d) Nb-D11 sensorgram and (e) response curve showing Nb-D11 has a high nanomolar affinity for Pf4 (K_d_ = 650 ± 60 nM).

**Extended Data Figure 2.**
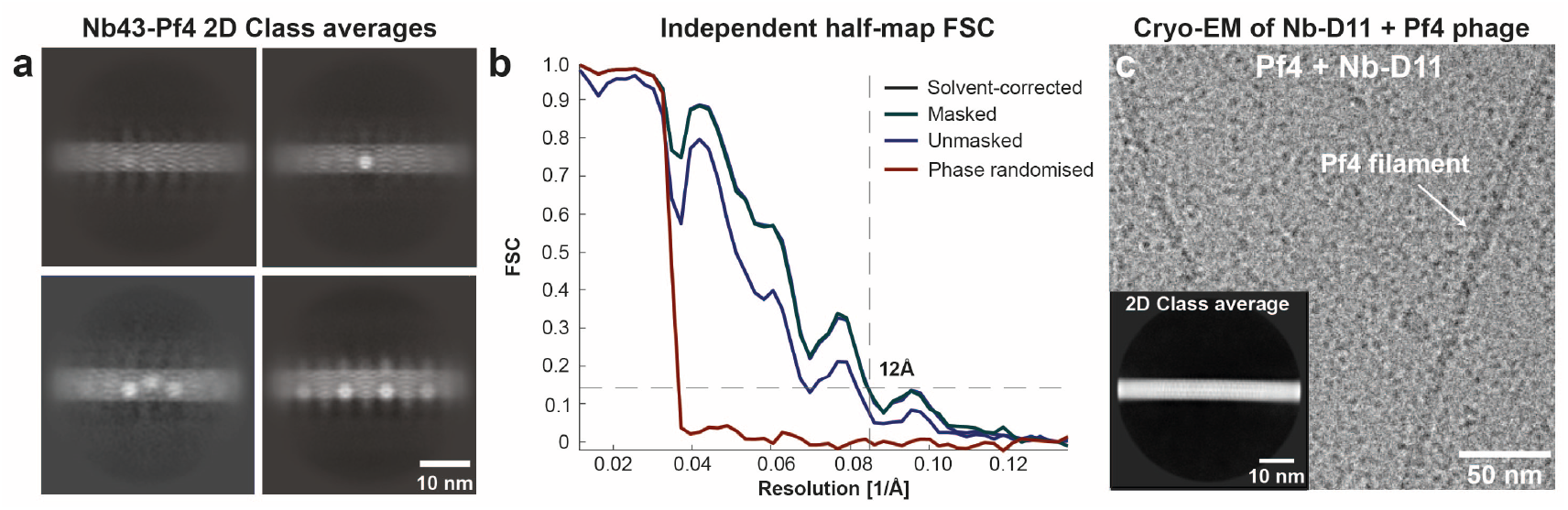
Cryo-EM analysis of Pf4 filaments incubated with nanobodies. (**a**) Gallery of 2D class averages illustrating heterogeneity of Nb43 occupancy on Pf4 filaments. (**b**) Resolution estimation of Pf4-Nb43 reconstruction (Fig. 2g) by Fourier shell correlation (FSC) of independently aligned and averaged half-maps. Dashed line indicates 0.143 criterion. (**c**) Cryo-EM micrograph of Pf4 with Nb-D11, showing Pf4 filaments with no apparent decoration. Inset shows a representative 2D class average.

**Extended Data Figure 3.**
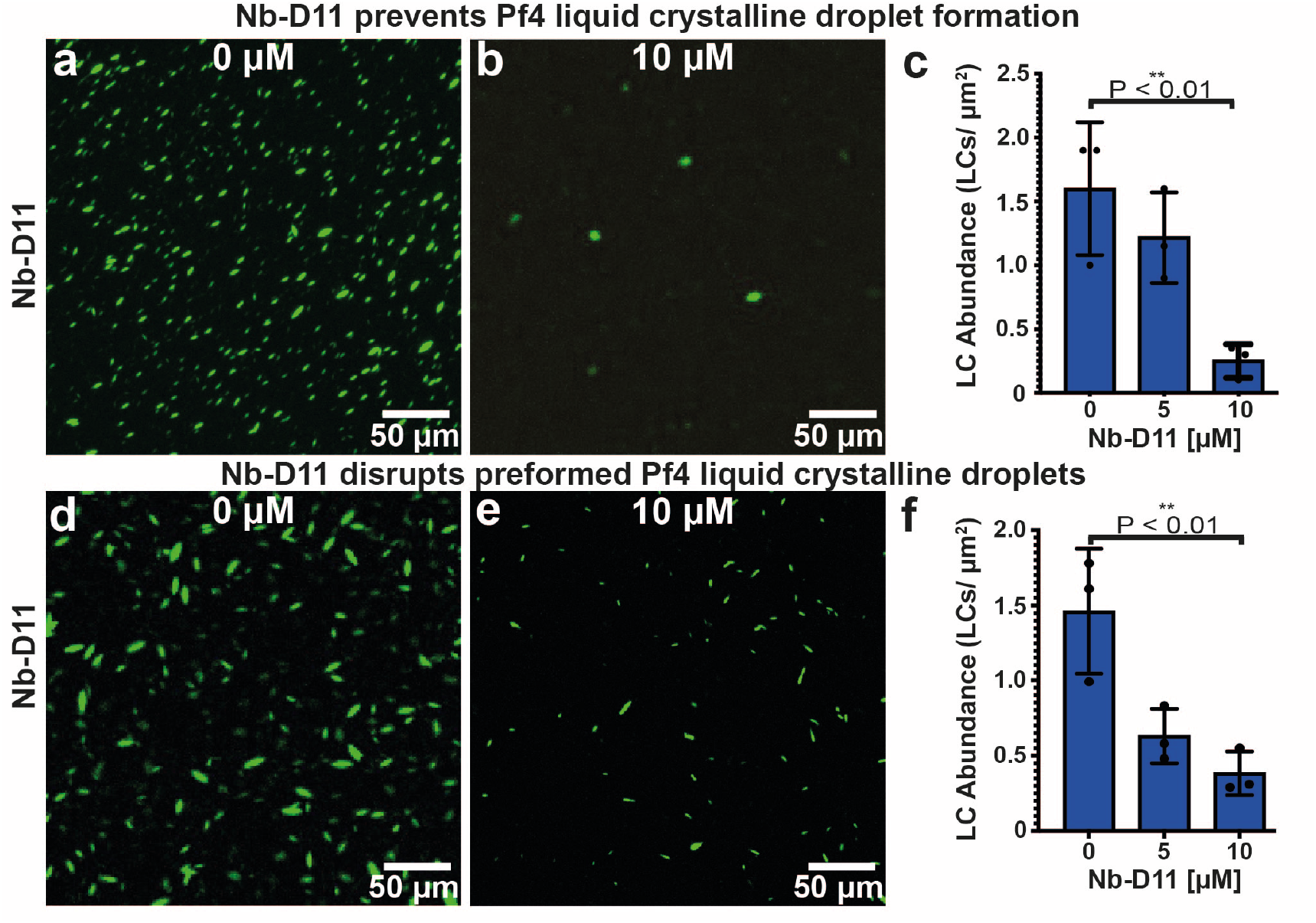
Nb-D11 prevents and disrupts Pf4 liquid crystalline droplets. (**a-b**) Representative light microscopy images of A488-labelled Pf4 liquid crystalline droplets formed in the presence of (a) no Nb-D11 or (b) 10 *μ*M Nb-D11. (**c**) Bar chart showing the abundance of Pf4 liquid crystalline droplets per *μ*m^2^. Addition of 10 *μ*M Nb-D11 results in a statistically significant reduction in liquid crystalline droplet formation (P_value_ < 0.01). (**d-e**) Representative light microscopy images of preformed liquid crystalline droplets treated with (d) no Nb-D11 or (e) 10 *μ*M Nb-D11. (**f**) Bar chart showing the abundance of Pf4 liquid crystalline droplets per *μ*m^2^ after treatment. Addition of 10 *μ*M Nb-D11 results in a statistically significant reduction in droplet abundance (P_value_ < 0.01). Error bars represent standard deviation. P-values were calculated using an unpaired t-test. All images have been background subtracted.

**Extended Data Figure 4.**
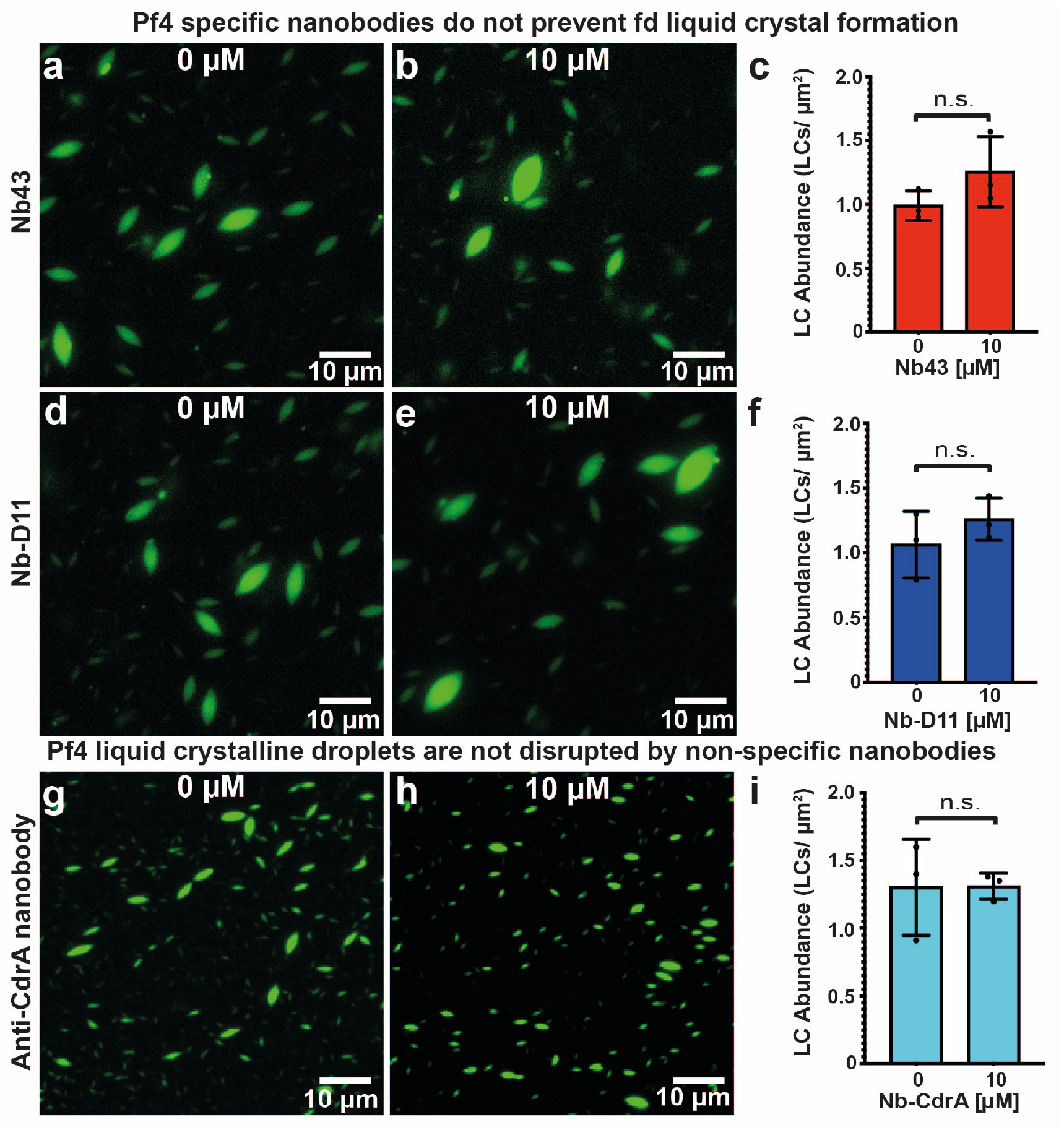
Disruption of Pf4 liquid crystalline droplets requires specific nanobody binding to Pf4 filaments. (**a-b**) Representative light microscopy images of A488-labelled fd liquid crystals formed in the presence of (a) no Nb43 or (b) 10 *μ*M Nb43. (**c**) Bar chart showing the abundance of fd liquid crystals per *μ*m2. Addition of 10 *μ*M Nb43 did not result in a statistically significant change in liquid crystal formation. (**d-e**) Representative light microscopy images of A488-labelled fd liquid crystals formed in the presence of (d) no Nb-D11 or (e) 10 *μ*M Nb-D11. (**f**) Bar chart showing the abundance of fd liquid crystals per *μ*m^2^. Addition of 10 *μ*M Nb-D11 did not result in a statistically significant change in liquid crystal formation. (**g-h**) Representative light microscopy images of A488-labelled Pf4 liquid crystalline droplets formed in the presence of (g) no nanobody or (h) 10 *μ*M anti-CdrA nanobody. (**i**) Bar chart showing the abundance of Pf4 liquid crystalline droplets per *μ*m^2^. Addition of 10 *μ*M anti-CdrA nanobody did not result in a statistically significant change in liquid crystalline droplet formation. All values are representative of 30 images from three independent replicates. Error bars represent standard deviation. P-values were calculated using an unpaired t-test. All images have been background subtracted.

**Extended Data Figure 5.**
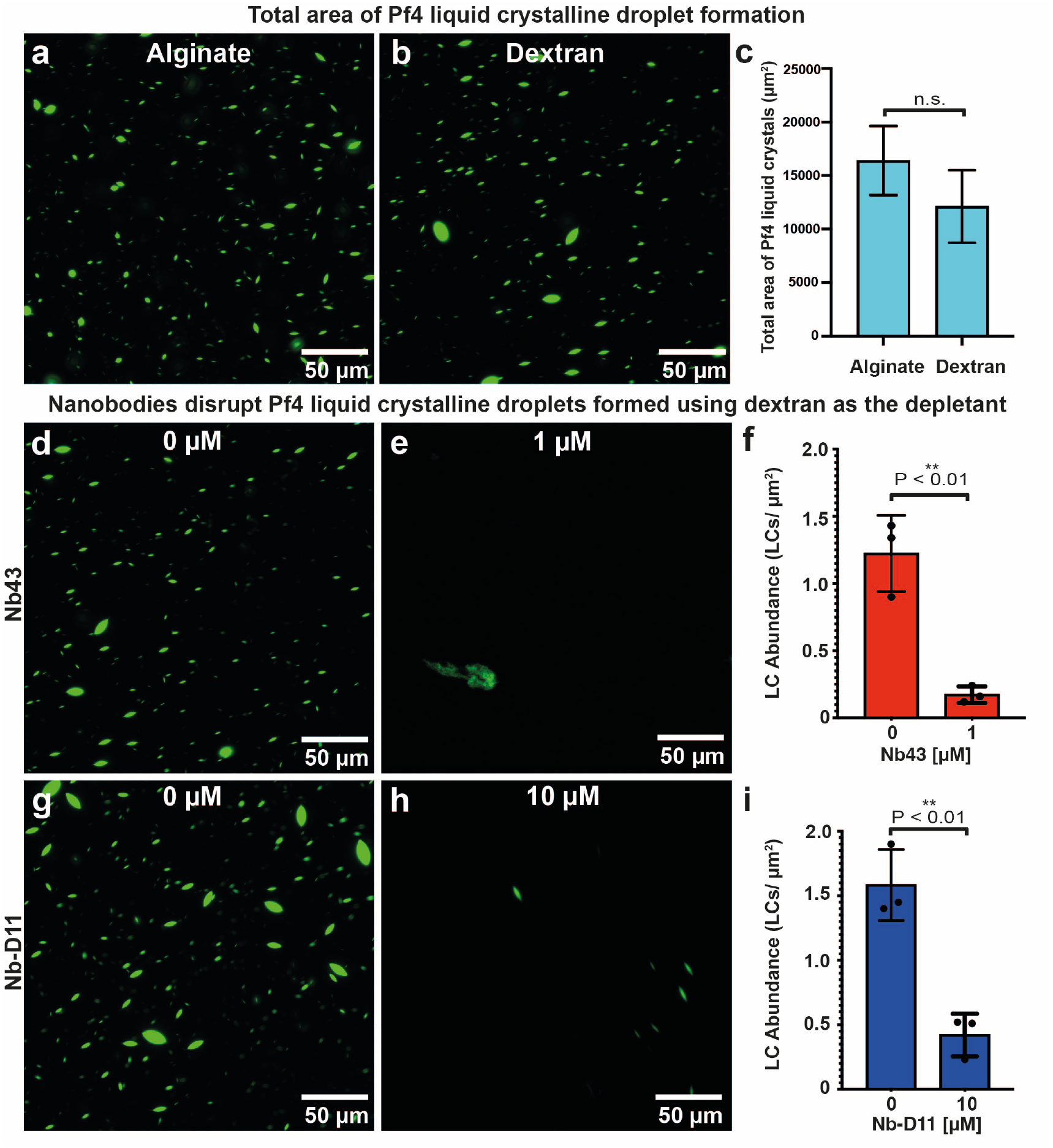
Nanobodies disrupt Pf4 liquid crystalline droplet formation using dextran as an alternative crowding agent (depletant). (**a-b**) Representative light microscopy images with comparable total liquid crystalline droplet area formed by A488-labelled Pf4 using (a) alginate and (b) dextran as the depletant. (**c**) Bar chart showing total liquid crystalline droplet area as assessed by light microscopy followed by image segmentation of droplets. (**d-e**) Representative light microscopy images of A488-labelled Pf4 liquid crystalline droplets formed with dextran as the depletant in the presence of (d) no Nb43 or (e) 1 *μ*M Nb43. (**f**) Bar chart showing the abundance of Pf4 liquid crystalline droplets per *μ*m2. Addition of 1 *μ*M Nb43 results in a statistically significant reduction in liquid crystalline droplet formation (P_value_ < 0.01). (**g-h**) Representative light microscopy images of A488-labelled Pf4 liquid crystalline droplets formed with dextran as the depletant in the presence of (g) no Nb-D11 or (h) 10 *μ*M Nb-D11. (**i**) Bar chart showing the abundance of Pf4 liquid crystalline droplets per *μ*m^2^. Addition of 10 *μ*M Nb-D11 results in a statistically significant reduction in liquid crystalline droplet formation (P_value_ < 0.01).

**Extended Data Figure 6.**
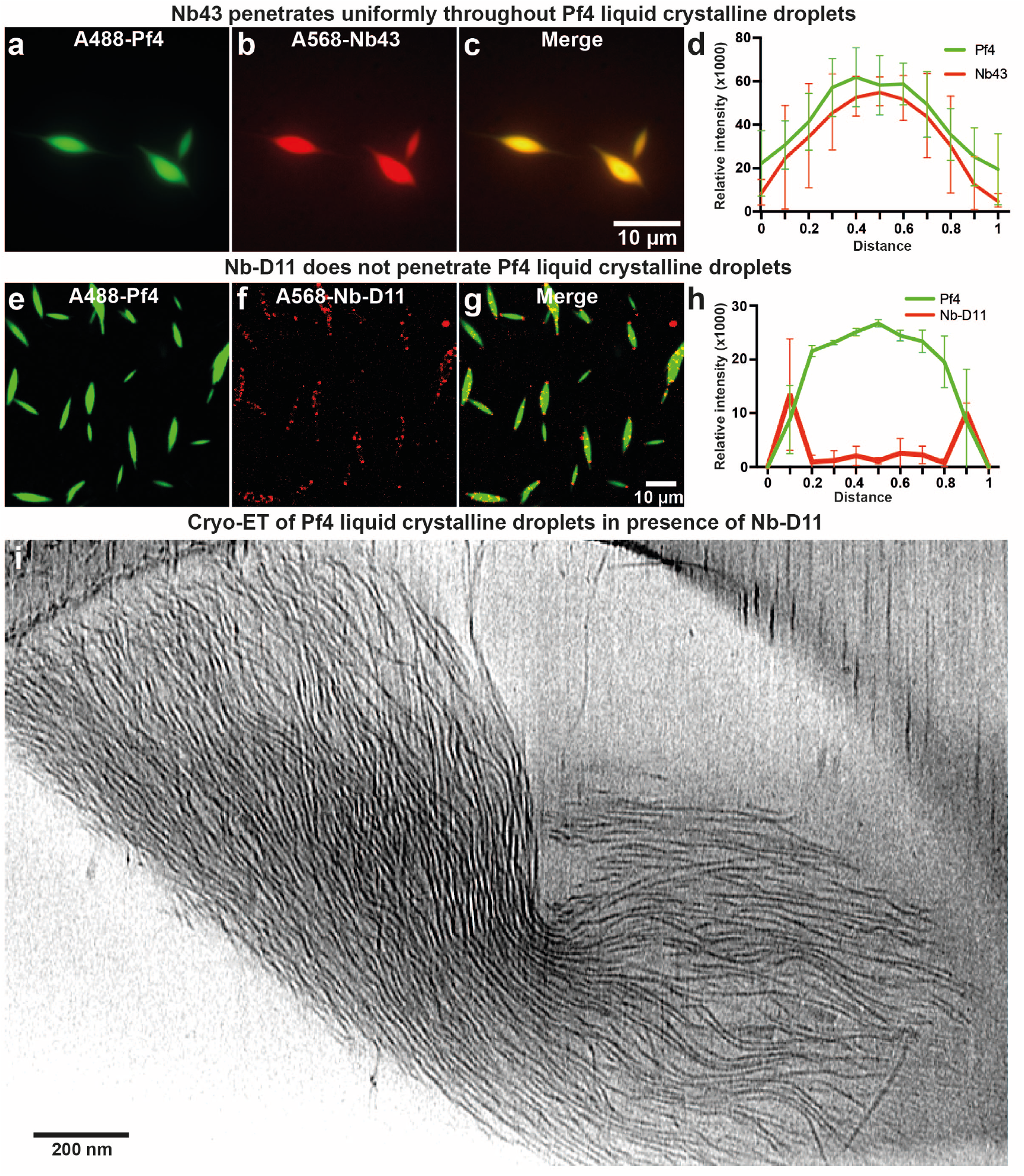
Nanobody binder localization in Pf4 liquid crystalline droplets. (**a-c**) Representative light microscopy images of A488-labelled Pf4 incubated with alginate and 0.1 *μ*M A568-labelled Nb43, (a) A488-Pf4 liquid crystalline droplets, (b) A568-labelled Nb43 and (c) merge. Images have been background subtracted. (**d**) Fluorescent intensity line profile through cross-section of Pf4 liquid crystalline droplet showing homogeneous signal for Nb43 (red line) throughout liquid crystalline droplets (green line). (**e-g**) Representative light microscopy images of A488-labelled Pf4 incubated with alginate and 2.5 *μ*M A568-labelled Nb-D11. Images have been background subtracted. (**h**) Fluorescent intensity line profile through cross-section of Pf4 liquid crystalline droplets showing signal for Nb-D11 (red line) is excluded to the tips and surface of the droplet (green line). (**i**) Cryo-ET slice of a Pf4 liquid crystalline droplet incubated with 2.5 *μ*M Nb-D11.

**Extended Data Figure 7.**
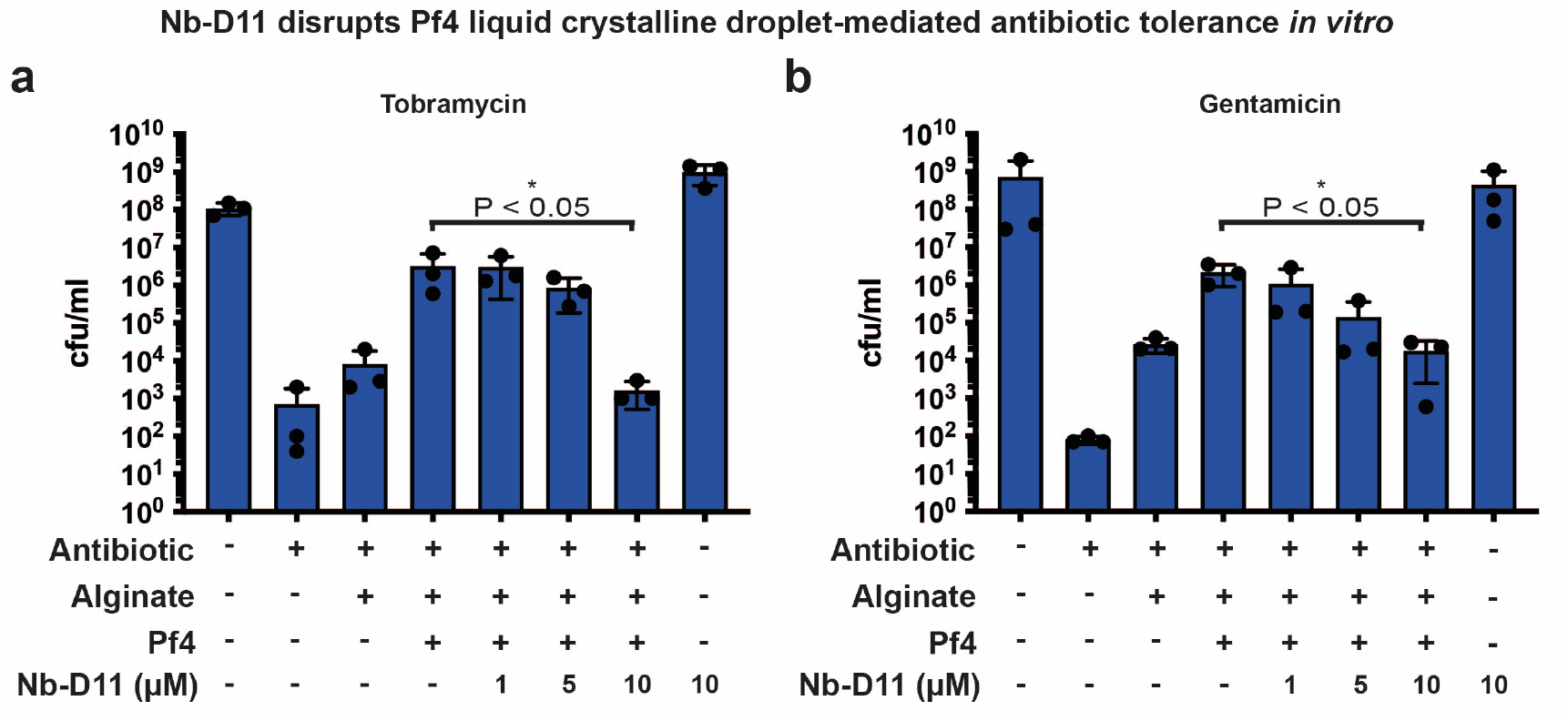
Nb-D11 disrupts phage liquid crystalline droplet-mediated antibiotic tolerance *in vitro* at high concentrations. (**a-b**) Bar graph shows colony-forming units (cfu) per ml (y axis), a measure of *P. aeruginosa* culture cell viability after (a) tobramycin and (b) gentamicin treatment in the presence of different reagents (x axis). For both antibiotics, 10 *μ*M Nb-D11 significantly reduced antibiotic tolerance levels to that seen in the alginate alone condition (P_value_ < 0.05). Values shown are the mean of three independent experiments. Error bars represent standard deviation. P-values were calculated using an unpaired t-test.

**Extended Data Figure 8.**
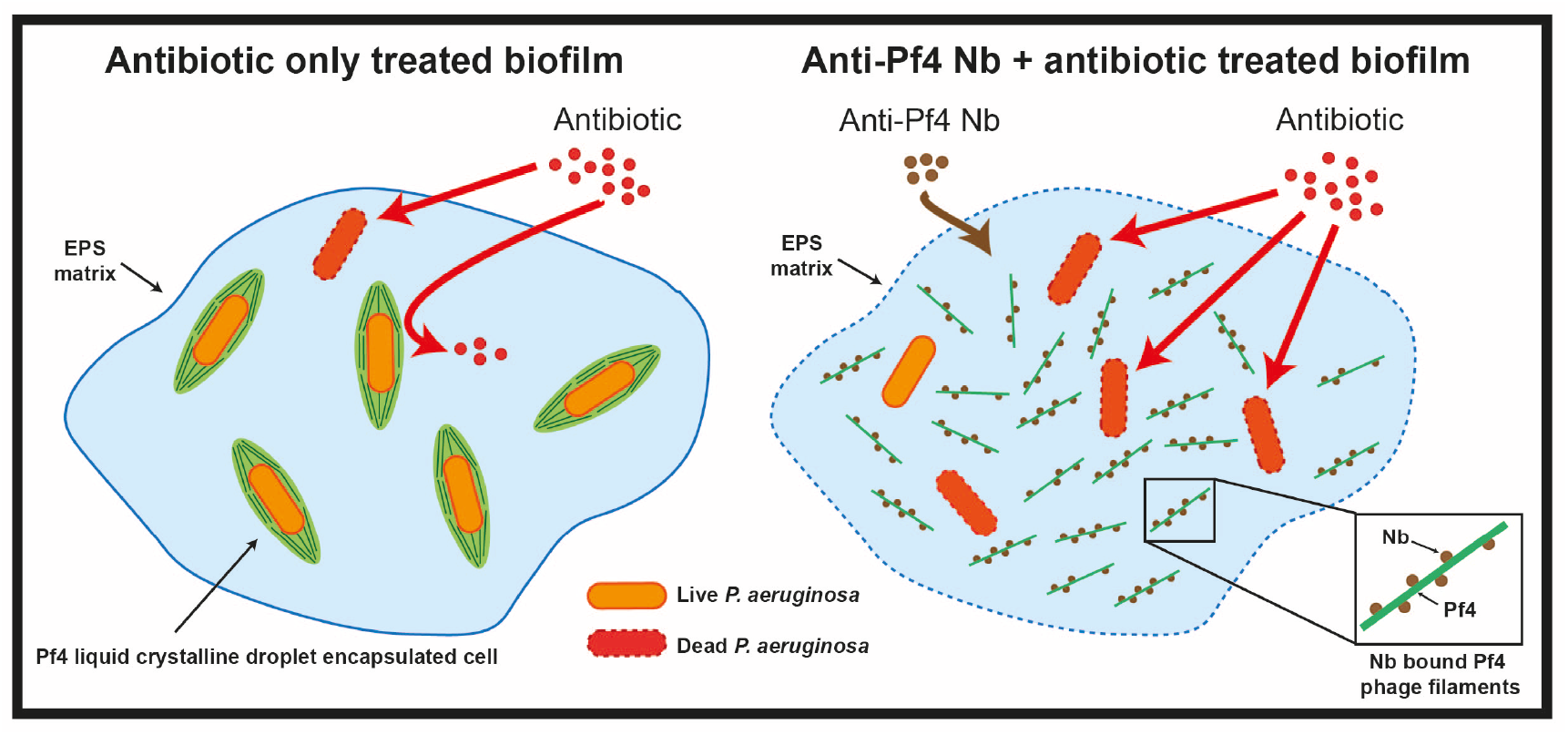
Schematic representation of nanobody action in abolishing antibiotic tolerance of *P. aeruginosa* biofilms. In untreated biofilms (left), cells show increased antibiotic tolerance due to Pf4 liquid crystalline droplets formed by depletion attraction in the biofilm EPS matrix, where encapsulated cells are protected by an antibiotic diffusion block. In nanobody treated biofilms, patchy binding of nanobody to Pf4 filaments reduces depletion attraction between the filaments preventing liquid crystalline droplet formation and encapsulation of cells, leading to increased antibiotic susceptibility of bacteria within the biofilm.

## Supplementary Video Legends

**Video S1. Tomogram of a Pf4 droplet treated with 0.1 *μ*M Nb43.** Sequential Z-slices of a tomogram of the sample of Pf4 liquid crystalline droplets treated with 0.1 *μ*M Nb43.

**Video S2. Tomogram of a Pf4 droplet treated with 1 *μ*M Nb43.** Sequential Z-slices of a tomogram of the sample of Pf4 liquid crystalline droplets treated with 1 *μ*M Nb43

## References

1. Flemming, H. C. & Wingender, J. The biofilm matrix. Nat Rev Microbiol 8, 623–633 (2010). 10.1038/nrmicro2415

2. Nadell, C. D., Drescher, K. & Foster, K. R. Spatial structure, cooperation and competition in biofilms. Nat Rev Microbiol 14, 589–600 (2016). 10.1038/nrmicro.2016.84

3. Stoodley, P., Sauer, K., Davies, D. G. & Costerton, J. W. Biofilms as complex differentiated communities. Annu Rev Microbiol 56, 187–209 (2002). 10.1146/annurev.micro.56.012302.160705

4. Böhning, J., Tarafder, A. K. & Bharat, T. A. M. The role of filamentous matrix molecules in shaping the architecture and emergent properties of bacterial biofilms. Biochem J 481, 245–263 (2024). 10.1042/BCJ20210301

5. Flemming, H. C. et al. Biofilms: an emergent form of bacterial life. Nat Rev Microbiol 14, 563–575 (2016). 10.1038/nrmicro.2016.94

6. Hoiby, N., Bjarnsholt, T., Givskov, M., Molin, S. & Ciofu, O. Antibiotic resistance of bacterial biofilms. Int J Antimicrob Agents 35, 322–332 (2010). 10.1016/j.ijantimicag.2009.12.011

7. Bouza, E., Burillo, A. & Munoz, P. Catheter-related infections: diagnosis and intravascular treatment. Clin Microbiol Infect 8, 265–274 (2002). 10.1046/j.1469-0691.2002.00385.x

8. Singh, P. K. et al. Quorum-sensing signals indicate that cystic fibrosis lungs are infected with bacterial biofilms. Nature 407, 762–764 (2000). 10.1038/35037627

9. Secor, P. R. et al. Filamentous bacteriophage promote biofilm assembly and function. Cell Host Microbe 18, 549–559 (2015). 10.1016/j.chom.2015.10.013

10. Tarafder, A. K. et al. Phage liquid crystalline droplets form occlusive sheaths that encapsulate and protect infectious rodshaped bacteria. Proc Natl Acad Sci U S A 117, 4724–4731 (2020). 10.1073/pnas.1917726117

11. Collaborators, G. B. D. A. R. Global burden of bacterial antimicrobial resistance 1990-2021: a systematic analysis with forecasts to 2050. Lancet 404, 1199–1226 (2024). 10.1016/S0140-6736(24)01867-1

12. Deusenbery, C., Wang, Y. & Shukla, A. Recent innovations in bacterial infection detection and treatment. ACS Infect Dis 7, 695–720 (2021). 10.1021/acsinfecdis.0c00890

13. Antimicrobial Resistance, C. Global burden of bacterial antimicrobial resistance in 2019: a systematic analysis. Lancet 399, 629–655 (2022). 10.1016/S0140-6736(21)02724-0

14. Valderrey, A. D. et al. Chronic colonization by Pseudomonas aeruginosa of patients with obstructive lung diseases: cystic fibrosis, bronchiectasis, and chronic obstructive pulmonary disease. Diagn Microbiol Infect Dis 68, 20–27 (2010). 10.1016/j.diagmicrobio.2010.04.008

15. Thi, M. T. T., Wibowo, D. & Rehm, B. H. A. Pseudomonas aeruginosa Biofilms. Int J Mol Sci 21 (2020). 10.3390/ijms21228671

16. Costerton, J. W. et al. Bacterial biofilms in nature and disease. Annu Rev Microbiol 41, 435–464 (1987). 10.1146/annurev.mi.41.100187.002251

17. O’Toole, G., Kaplan, H. B. & Kolter, R. Biofilm formation as microbial development. Annu Rev Microbiol 54, 49–79 (2000). 10.1146/annurev.micro.54.1.49

18. Hall-Stoodley, L. & Stoodley, P. Evolving concepts in biofilm infections. Cell Microbiol 11, 1034–1043 (2009). 10.1111/j.1462-5822.2009.01323.x

19. Ceri, H. et al. The Calgary Biofilm Device: new technology for rapid determination of antibiotic susceptibilities of bacterial biofilms. J Clin Microbiol 37, 1771–1776 (1999). 10.1128/JCM.37.6.1771-1776.1999

20. Rice, S. A. et al. The biofilm life cycle and virulence of Pseudomonas aeruginosa are dependent on a filamentous prophage. ISME J 3, 271–282 (2009). 10.1038/ismej.2008.109

21. Webb, J. S. et al. Cell death in Pseudomonas aeruginosa biofilm development. J Bacteriol 185, 4585–4592 (2003). 10.1128/JB.185.15.4585-4592.2003

22. Marvin, D. A., Symmons, M. F. & Straus, S. K. Structure and assembly of filamentous bacteriophages. Prog Biophys Mol Biol 114, 80–122 (2014). 10.1016/j.pbiomolbio.2014.02.003

23. Whiteley, M. et al. Gene expression in Pseudomonas aeruginosa biofilms. Nature 413, 860–864 (2001). 10.1038/35101627

24. Secor, P. R. et al. Pf bacteriophage and their impact on Pseudomonas virulence, mammalian immunity, and chronic infections. Front Immunol 11, 244 (2020). 10.3389/fimmu.2020.00244

25. Burgener, E. B. et al. Filamentous bacteriophages are associated with chronic Pseudomonas lung infections and antibiotic resistance in cystic fibrosis. Sci Transl Med 11 (2019). 10.1126/scitranslmed.aau9748

26. Sweere, J. M. et al. Bacteriophage trigger antiviral immunity and prevent clearance of bacterial infection. Science 363 (2019). 10.1126/science.aat9691

27. Secor, P. R. et al. Filamentous bacteriophage produced by Pseudomonas aeruginosa alters the inflammatory response and promotes noninvasive infection in vivo. Infect Immun 85 (2017). 10.1128/IAI.00648-16

28. Bach, M. S. et al. Filamentous bacteriophage delays healing of Pseudomonas-infected wounds. Cell Rep Med 3, 100656 (2022). 10.1016/j.xcrm.2022.100656

29. Lekkerkerker, H. N. & Tuinier, R. in Colloids and the Depletion Interaction 57–108 (Springer, 2011).

30. Böhning, J. et al. Biophysical basis of filamentous phage tactoid-mediated antibiotic tolerance in P. aeruginosa. Nat Commun 14, 8429 (2023). 10.1038/s41467-023-44160-8

31. Böhning, J. et al. Structure of the virulence-associated Neisseria meningitidis filamentous bacteriophage MDAF. bioRxiv, 2024.2010.2002.616247 (2024). 10.1101/2024.10.02.616247

32. Waldor, M. K. & Mekalanos, J. J. Lysogenic conversion by a filamentous phage encoding cholera toxin. Science 272, 1910–1914 (1996). 10.1126/science.272.5270.1910

33. Karygianni, L., Ren, Z., Koo, H. & Thurnheer, T. Biofilm matrixome: extracellular components in structured microbial communities. Trends Microbiol 28, 668–681 (2020). 10.1016/j.tim.2020.03.016

34. Chapman, M. R. et al. Role of Escherichia coli curli operons in directing amyloid fiber formation. Science 295, 851–855 (2002). 10.1126/science.1067484

35. Böhning, J. et al. Donor-strand exchange drives assembly of the TasA scaffold in Bacillus subtilis biofilms. Nat Commun 13, 7082 (2022). 10.1038/s41467-022-34700-z

36. Fernandez-Martinez, D. et al. Cryo-EM structures of type IV pili complexed with nanobodies reveal immune escape mechanisms. Nat Commun 15, 2414 (2024). 10.1038/s41467-024-46677-y

37. Melia, C. E. et al. Architecture of cell-cell junctions in situ reveals a mechanism for bacterial biofilm inhibition. Proc Natl Acad Sci U S A 118 (2021). 10.1073/pnas.2109940118

38. Adams, H. et al. Inhibition of biofilm formation by Camelid single-domain antibodies against the flagellum of Pseudomonas aeruginosa. J Biotechnol 186, 66–73 (2014). 10.1016/j.jbiotec.2014.06.029

39. Maksymova, L., Pilger, Y. A., Nuhn, L. & Van Ginderachter, J. A. Nanobodies targeting the tumor microenvironment and their formulation as nanomedicines. Mol Cancer 24, 65 (2025). 10.1186/s12943-025-02270-5

40. Zhao, K. & Mason, T. G. Suppressing and enhancing depletion attractions between surfaces roughened by asperities. Phys Rev Lett 101, 148301 (2008). 10.1103/PhysRevLett.101.148301

41. Badaire, S., Cottin-Bizonne, C., Woody, J. W., Yang, A. & Stroock, A. D. Shape selectivity in the assembly of lithographically designed colloidal particles. J Am Chem Soc 129, 40–41 (2007). 10.1021/ja067527h

42. Badaire, S., Cottin-Bizonne, C. & Stroock, A. D. Experimental investigation of selective colloidal interactions controlled by shape, surface roughness, and steric layers. Langmuir 24, 11451–11463 (2008). 10.1021/la801718j

43. Zhao, K. & Mason, T. G. Directing colloidal self-assembly through roughness-controlled depletion attractions. Phys Rev Lett 99, 268301 (2007). 10.1103/PhysRevLett.99.268301

44. Barry, E. & Dogic, Z. Entropy driven self-assembly of nonamphiphilic colloidal membranes. Proc Natl Acad Sci U S A 107, 10348–10353 (2010). 10.1073/pnas.1000406107

45. Kamp, M. et al. Selective depletion interactions in mixtures of rough and smooth silica spheres. Langmuir 32, 1233–1240 (2016). 10.1021/acs.langmuir.5b04001

46. Lopes, M. A. et al. Probing insulin bioactivity in oral nanoparticles produced by ultrasonication-assisted emulsification/internal gelation. Int J Nanomedicine 10, 5865–5880 (2015). 10.2147/IJN.S86313

47. Zupancic, J. M. et al. Directed evolution of potent neutralizing nanobodies against SARS-CoV-2 using CDR-swapping mutagenesis. Cell Chem Biol 28, 1379–1388 e1377 (2021). 10.1016/j.chembiol.2021.05.019

48. Whitchurch, C. B., Tolker-Nielsen, T., Ragas, P. C. & Mattick, J.S. Extracellular DNA required for bacterial biofilm formation. Science 295, 1487 (2002). 10.1126/science.295.5559.1487

49. DiGiandomenico, A. et al. A multifunctional bispecific antibody protects against Pseudomonas aeruginosa. Sci Transl Med 6, 262ra155 (2014). 10.1126/scitranslmed.3009655

50. Bille, E. et al. A virulence-associated filamentous bacteriophage of Neisseria meningitidis increases host-cell colonisation. PLoS Pathog 13, e1006495 (2017). 10.1371/journal.ppat.1006495

51. Thompson, A. P. et al. LAMMPS-a flexible simulation tool for particle-based materials modeling at the atomic, meso, and continuum scales. Computer physics communications 271, 108171 (2022).

52. Collins, A. J., Pastora, A. B., Smith, T. J. & O’Toole, G. A. MapA, a second large RTX adhesin conserved across the Pseudomonads, contributes to biofilm formation by Pseudomonas fluorescens. J Bacteriol 202 (2020). 10.1128/JB.00277-20

53. Pardon, E. et al. A general protocol for the generation of nanobodies for structural biology. Nat Protoc 9, 674–693 (2014). 10.1038/nprot.2014.039

54. Eyssen, L. E. et al. From Llama to nanobody: a streamlined workflow for the generation of functionalised VHHs. Bio Protoc 14, e4962 (2024). 10.21769/BioProtoc.4962

55. Mastronarde, D. N. Automated electron microscope tomography using robust prediction of specimen movements. J Struct Biol 152, 36–51 (2005). 10.1016/j.jsb.2005.07.007

56. Burt, A. et al. An image processing pipeline for electron cryo-tomography in RELION-5. FEBS Open Bio 14, 1788–1804 (2024). 10.1002/2211-5463.13873

57. Zheng, S. et al. AreTomo: An integrated software package for automated marker-free, motion-corrected cryo-electron tomographic alignment and reconstruction. J Struct Biol X 6, 100068 (2022). 10.1016/j.yjsbx.2022.100068

58. Buchholz, T.-O., Jordan, M., Pigino, G. & Jug, F. in 2019 IEEE 16th International Symposium on Biomedical Imaging (ISBI 2019). 502–506.

59. Buchholz, T.-O. et al. in Methods in Cell Biology Vol. 152 Three-Dimensional Electron Microscopy (eds Thomas Müller-Reichert & Gaia Pigino) 277–289 (Academic Press, 2019).

60. Punjani, A., Rubinstein, J. L., Fleet, D. J. & Brubaker, M. A. cryoSPARC: algorithms for rapid unsupervised cryo-EM structure determination. Nat Methods 14, 290–296 (2017). 10.1038/nmeth.4169

61. Ducret, A., Quardokus, E. M. & Brun, Y. V. MicrobeJ, a tool for high throughput bacterial cell detection and quantitative analysis. Nat Microbiol 1, 16077 (2016). 10.1038/nmicrobiol.2016.77

62. Schindelin, J. et al. Fiji: an open-source platform for biologicalimage analysis. Nat Methods 9, 676–682 (2012). 10.1038/nmeth.2019

63. Chan, T. F. & Vese, L. A. Active contours without edges. IEEE Trans Image Process 10, 266–277 (2001). 10.1109/83.902291

64. O’Toole, G.A. Microtiter dish biofilm formation assay. J Vis Exp (2011). 10.3791/2437

65. McDonald, J. C. et al. Fabrication of microfluidic systems in poly(dimethylsiloxane). Electrophoresis 21, 27–40 (2000). 10.1002/(SICI)1522-2683(20000101)21:1

66. Drescher, K. et al. Architectural transitions in Vibrio cholerae biofilms at single-cell resolution. Proc Natl Acad Sci U S A 113, E2066–2072 (2016). 10.1073/pnas.1601702113

67. Shcherbo, D. et al. Bright far-red fluorescent protein for wholebody imaging. Nat Methods 4, 741–746 (2007). 10.1038/nmeth1083

